# Bead-Ejection Scenario in Electrospray Ionization of Multi-Domain Nucleic Acids

**DOI:** 10.1101/2025.07.29.667413

**Authors:** Debasmita Ghosh, Frédéric Rosu, Valérie Gabelica

**Affiliations:** Univ. Bordeaux, INSERM, CNRS, ARNA, UMR 5320, U1212, F-33000 Bordeaux, France; Univ. Bordeaux, CNRS, INSERM, IECB, UAR3033, US01, F-33600 Pessac, France; School of Pharmaceutical Sciences, University of Geneva, Geneva, Switzerland

## Abstract

Understanding how biomolecules acquire their charge and retain their solution conformation during electrospray ionization (ESI) is crucial for native mass spectrometry (native MS) interpretation. Here, we examine the charging and gas phase conformation of nucleic acid constructs comprising folded G-quadruplex “beads” linked by unstructured polythymine regions. Under physiological ionic strength, these oligonucleotides exhibit a multimodal charge-state and collision cross section distribution, revealing multiple conformational ensembles, in contrast to the unimodal profiles typically observed for shorter oligonucleotides. Native MS observations for intermediate charge states are compatible with ion production via the recently proposed bead ejection scenario, in addition to the charge residue scenario for low charge states and chain ejection for the highest charge states or for sequences with thymine overhangs on both ends. The preservation of the local structures in ions charged above the Rayleigh limit helps to infer the presence of folded subunits. The position of the G-quadruplex subunit and ionic strength governs the charging and retention of G-quadruplex folded regions. Our findings broaden the existing conceptual framework underpinning nucleic acid ionization.

**TOC graphic:** 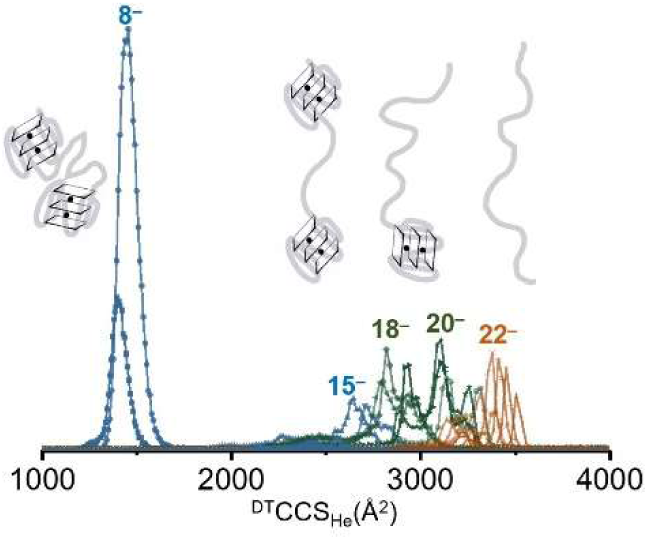

## Introduction

Native electrospray ionization (ESI) mass spectrometry (MS) preserves the covalent and non-covalent interactions of biomolecules during their transition from solution to the gas phase.^1^ Although ESI was introduced over three decades ago,^2^ the mechanisms that govern the final stages of ion charging and desolvation remain a matter of debate. Three major models are commonly discussed. In the ion evaporation model (IEM),^3^ field emission of ions can occur directly from the surface of the droplet when the electrostatic repulsion exceeds the solvation energy. Although the IEM is primarily thought to apply to small ions^4, 5^ it was also observed in molecular dynamics simulations for folded proteins.^6^ In the charged residue model (CRM),^7^ the electrospray droplet evaporates to dryness around the analyte, which will carry a residual charge that cannot exceed the Rayleigh limit charge for a droplet of the ion size.^5, 8^ The CRM is thought to apply to large globular analytes.^9^ More recently, the chain ejection model (CEM)^10, 11^ was proposed for disordered solvophobic (bio)polymers, which progressively extrude from the droplet in extended conformations, carrying away numerous charges.

Although presenting these models as distinct mechanisms aids teaching, we believe such rigid categorization can lead to confusion. The IEM, CRM and CEM should be viewed more (1) as limiting cases, and (2) as “scenarios” (or pathways) rather than mechanisms, i.e. sequence of events, for which we experimentally observe only the ending: the charge state distribution (CSD) and the collision cross section distribution (CCSD) for each charge state. After all, the underlying physics is the same—repulsion between like charges, counterbalanced by non-covalent intramolecular forces (folding), analyte-solvent and solvent-solvent interaction forces—and only the balance between forces is different.

Moreover, the reality is sometimes more complex than assuming that one class of analyte will always be ionized via the same scenario. First, intermediate scenarios can occur.^12^ For example, we proposed that multi-domain proteins can ionize via a hybrid CRM-CEM route called the bead ejection model (BEM), wherein folded domains (beads) remain compact (local CRM) while the disordered linkers extend (local CEM).^13^ Second, within a single sample, different fractions of the analyte population may follow distinct ionization pathways, even though the solution-phase population is uniform. For example, in unstructured nucleic acid single strands sprayed in the negative ion mode,^14, 15^ we found that the ionic strength was the major factor affecting the proportion of low charge state compact ions (CRM) vs. high charge state extended ions (CEM). High ammonium acetate concentrations (100-150 mM, physiological ionic strength) favor the CRM scenario because the presence of more acetate anions (charge carriers) on the droplet surface allows the nucleic acids to stay in the droplet center. At the charge states resulting from this process, even collision-induced unfolding experiments fail to reveal the contribution of specific intramolecular folding, because the unfolding does not result in any significant extension (contraction is sometimes observed instead).^16^ In contrast, low [NH_4_OAc] favors the CEM scenario.^15, 17^ As a result, the CSD and CCSD of nucleic acids during ESI can depend more on the ionization process than on the solution folding state.^15^

Most of our previous work has dealt with short (20—30 base) oligonucleotides, folded or unfolded in solution, and the results could be rationalized by invoking the CRM and CEM scenarios. Here, we wanted to establish whether the BEM hybrid scenario could apply to longer nucleic acids in the negative ion mode. We used sequences containing G-quadruplexes as model systems. Folded G-quadruplex nucleic acid structures are stabilized by specific coordination of cations (NH_4_^+^, K^+^) within guanine quartets. Both the nature and concentration of these cations determine the structure and stability of G-quadruplexes.^18, 19^ Ammonium acetate is commonly used in native MS as a volatile electrolyte, with its concentration chosen based on the desired ionic strength.^20^ Inspired by our findings for multi-domain proteins,^13^ we designed mixed nucleic acid structures containing both G-quadruplex (G4) subunit and unstructured polythymine (Tn) sequences, placed at various positions (Table 1). As a control, we examined a non-folding single strand sequence (NG) which lacks enough contiguous G-tracts to form a G-quadruplex and thus has no specific NH_4_^+^ coordination sites. Each G4 subunit contains 3 G-quartets and thus 2 specifically bound NH_4_^+^ ions. Thus, the mass measurement informs us on how many G4 subunits are preserved in the gas phase. We also used native ion mobility MS^21–23^ to deduce the gas-phase compactness of the gas-phase structures and deduce the possible ion production scenarios that could lead to the different conformers.

**Table 1.** 60-mer DNA sequences studied herein.

| Name | Sequence | Number of Base | Structure |
| --- | --- | --- | --- |
| G4Tn | [dT(TG<sub>3</sub>)<sub>4</sub>T<sub>43</sub>•(NH<sub>4</sub><sup>+</sup>)<sub>2</sub>] | 60 | 1G4 |
| TnG4 | [dT<sub>43</sub>T(TG<sub>3</sub>)<sub>4</sub>•(NH<sub>4</sub><sup>+</sup>)<sub>2</sub>] | 60 | 1G4 |
| TnG4Tn | [dT<sub>21</sub>T(TG<sub>3</sub>)<sub>4</sub>T<sub>22</sub>•(NH<sub>4</sub><sup>+</sup>)<sub>2</sub>] | 60 | 1G4 |
| G4TnG4 | [dT(TG<sub>3</sub>)<sub>4</sub>T<sub>26</sub>T(TG<sub>3</sub>)<sub>4</sub>•(NH<sub>4</sub><sup>+</sup>)<sub>4</sub>] | 60 | 2G4 |
| NG | [dT<sub>4</sub>GT<sub>5</sub>TGT<sub>4</sub>(GT<sub>3</sub>)<sub>5</sub>(GT<sub>2</sub>)<sub>2</sub>GT<sub>4</sub>TGT<sub>4</sub>GT<sub>4</sub>T<sub>2</sub>] | 60 | non-structured |

### Experimental Section

#### Sample Preparation

All single strands were purchased from Eurogentec (Seraing, Belgium, with RP cartridge purification) and dissolved in water from Biosolve. Desalting is crucial for our sample preparation, particularly since some experiments are conducted at low ionic strength. We used Amicron ultracel-3 centrifugal filters (Merck Millipore Ltd.) with a 3-kDa molecular weight cutoff. The samples underwent several centrifugation passes with a 200 mM aqueous NH_4_OAc solution, followed by several passes with a 100 mM aqueous NH_4_OAc solution, and finally with water to achieve satisfactory desalting.

All G-quadruplexes were formed in 150 mM ammonium acetate. Intramolecular G-quadruplexes were formed from 200 μM single strand, incubated for at least 24 h in the folding buffer (150 mM aqueous NH_4_OAc). Single strands were annealed at 85°C in water for 5 min to ensure the removal of any preformed structures. Control experiments were done using non-structured oligonucleotides of the same length. The supercharging agent propylene carbonate (PC) (99.7% purity) was purchased from Sigma-Aldrich. The DNA structures were diluted and injected at a final concentration of 15 μM final concentration for IM-MS analysis.

#### UV spectroscopy

The absorbance was recorded at 260 nm using a Uvikon XS spectrophotometer and the stock sample concentrations were determined using the Beer−Lambert law. Molar extinction coefficients were estimated using the nearest-neighbor model and Cantor’s parameters.^24^ The stability of G4 structures in solution was examined with a UVmc2 double-beam spectrophotometer (SAFAS, Monte Carlo, Monaco) equipped with a temperature-controlled 10-cell holder and a high-performance Peltier temperature controller.

Thermal denaturation was monitored by measuring the changes in the absorbance at 295 nm as a function of temperature. Samples were heated to 90 °C, then the absorbance was recorded at 260, 295, and 350 nm in a series contains a cooling down to 4 °C at a rate of 0.2 °C min^−1^, followed by reheating to 90 °C at the same rate.

The resulting absorbance versus temperature data were converted into a fraction folded (θ) versus temperature plot.^25^ We fitted the upper and lower baselines, then calculated the fraction folded at each temperature, *θ*_T_ (Eq. 1), where L0_T_ and L1_T_ are baseline values of the unfolded and folded species. The temperature at which θ = 0.5 is known as the melting temperature.^25^

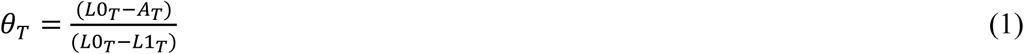

#### Native ion mobility mass spectrometry (IM-MS)

IM-MS and collision-induced unfolding (CIU) experiments were conducted at 24 °C using a 6560 DTIMS-Q-TOF instrument (Agilent Technologies, Santa Clara, CA) adapted to run with helium in the drift tube.^26^ All experiments were performed in negative ion mode with the standard electrospray ionization (ESI) source. A syringe pump flow rate of 3 µL/min was used. The parameters used were fragmentor = 320 V (unless stated otherwise for CIU experiments), nebulizing gas = 4 psi, drying gas = 1 L/min, trap fill time = 1000 μs, trap release time = 150 μs, and trap entrance grid delta (TEGD) = 2 V. For collision cross-section (CCS) quality control, [(dTG_4_T)_4_(NH_4_)_3_]^5−^ was injected prior to the analysis, ensuring a ^DT^CCS_He_ within 785−791 Å² of its previously determined value (788 Å²).^27^ The data extraction was performed using the IM-MS Browser software version B.08.00 (Agilent Technologies).

The conversion of arrival time distributions to CCS distributions is performed as follows.^28^ First, the CCS of the centroid of one of the peaks is measured using the step-field method: five IM-MS spectra (segments) are recorded with varying drift tube entrance voltages (650, 700, 800, 900, 1000 V), and the arrival time of the centroid of the peak (*t*_A_) is measured as a function of the voltage difference between the entrance and exit of the drift tube (ΔV). A linear regression with Eq. (2) provides CCS from the slope, knowing that (*μ* = *m*_gas_*m*_ion_/(*m*_gas_ + *m*_ion_), *p*_0_ is the standard pressure (760 Torr), *p* is the pressure in the drift tube (3.89 ± 0.01 Torr), *T*_0_ is the standard temperature (273.15 K), *T* is the temperature in the tube (296.15 ± 1 K), *N*_0_ is the buffer gas number density at *T*_0_ and *p*_0_ (2.687 10^25^ m^−3^), *L* is the physical length of the drift tube (78.1 ± 0.2 cm), *k*_B_ is the Boltzmann constant, *z* is the absolute value of the ion nominal charge, and *e* is the charge of the electron.

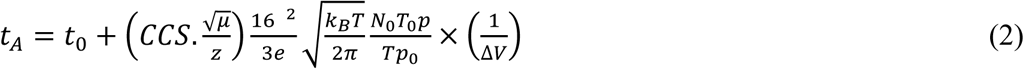

To reconstruct all CCS distributions from the arrival time distributions, the CCS determined with the step-field method and *t*_A_ determined for the 650 V segment are used to calculate the factor *a* using Equation (3), which is then used to change the axes from *t*_A_ (recorded at 650 V) to CCS for all other peaks.

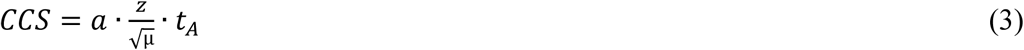

Collision-Induced Unfolding (CIU) experiments were conducted by increasing the fragmentor voltage from 300V to 450V. 20 V steps at 2-minute intervals were acquired. The arrival time distributions were plotted at different fragmentor voltages to compare the gas phase stabilities of 60-mers.

#### CCS calculations for gas-phase structural models

Structures of G_4_T_n_ were generated in vacuo using molecular dynamics simulations and the theoretical collision cross sections were calculated using EHSSrot^29^, with Siu’s modified atom size parameters in helium.^30^ The propeller-type parallel-stranded G-quadruplex 5’-TTGGGTGGGTGGGTGGGT-3’ is taken from the PDB code 2LK7^31^ and modified by coupling a 42-mer strand of thymine. The charge states (8- and 29-) were generated manually using localized charges (protons). Using a localized or delocalized charges model has only a small impact on the generated structure when using force fields (amber^32^ with parmbsc1^33^, using using Hyperchem 8.0.10 (Hypercube, Inc.)). The position of the charge has a important effect on the structure only when using higher level of calculation like semi-empirical PM7^34^ or DFT. Given the size of the oligonucleotide (60-mer), only force field is used on the complete sequence. PM7 will be used only on portions of the structure (the G-quadruplex structure, and two single strands truncated to dT_21_). The three parts are then concatenated to calculate the collision cross section for low charge states.

For the extended forms of G4Tn, we generated using Hyperchem a 42-mer of homothymine with regular B-helix and connected it to the quadruplex structure. Starting from the 3’end, 29 charges were located on the phosphate groups (i.e., one phosphate deprotonated follow by a neutralized one). Molecular dynamics was performed for 40 ns. The structure elongates rapidly (after less than 1 ns) to reach an extended structure. To simulate more compact forms of (G4Tn)^8-^, another structure of the homothymine strand dT_42_ has been generated using the curvature of the DNA on the histone H2 (pdb code: 7LV9).^35^ A 42-mer was extracted and the bases were mutated to thymine. The structure was cut into two 21-mers and optimized using Gaussian 16 rev. C01^36^ at the semi-empirical level of theory (PM7). Each dT_21_ carries three negative charges arbitrarily located every 10 bases. The G-quadruplex with 2- charge is also optimized at the PM7 level and concatenated to the thymine strands. TnG4Tn was generated with the concatenation of the homothymine strands at each extremities, and the same procedure as for G4Tn was applied. Finally, as these structures gave CCS values still larger than the experimentally observed ones for low charge states, we built a third model for G4Tn, using concatenations of dT_6_ strands that had been optimized at the DFT level of theory. dT6 forms intramolecular hydrogen bonds with the surrounding bases and with the phosphate backbones rendering it very compact. We previously demonstrated the compactness of the DNA single strands in-vacuo using ion mobility and resolved wavelength ion spectroscopy.^37^

## Results and Discussion

### 60-mer oligonucleotides in physiological ionic strength produce multimodal charge state distributions

To investigate ESI mechanisms in nucleic acid structures containing both folded and unfolded domains, we first examined 60-mer oligonucleotides with one G-quadruplex (G4) subunit: G4Tn, TnG4, and TnG4Tn. One parallel-stranded intramolecular G4 is stabilized by two specifically coordinated NH_4_^+^ ions,^18^ while polythymine chains are unstructured. The 60-mer single-stranded sequence NG is used as a control. The full sequences are listed in Table 1, and we confirmed that all G4-containing constructs are folded at room temperature (Figure S1).

Native electrospray MS spectra of these 60-mers obtained from 150 mM aqueous NH_4_OAc (Figure 1) show a low charge state distribution (mainly 7- and 8-), and a higher charge state distribution (from 14- to 22-, or even up to 26-, depending on the constructs). This finding contrasts with oligonucleotides up to 24-mers, which typically only exhibit low charges (such as 4- or 5-) at comparable ionic strength. Thus, although the G4 subunit is folded in solution, native MS data reveal multimodal CSDs at physiological ionic strength. We suspect that the unstructured polythymine tails drive the formation of high charge states, and this will be discussed further in the rest of the manuscript.

**Figure 1.**
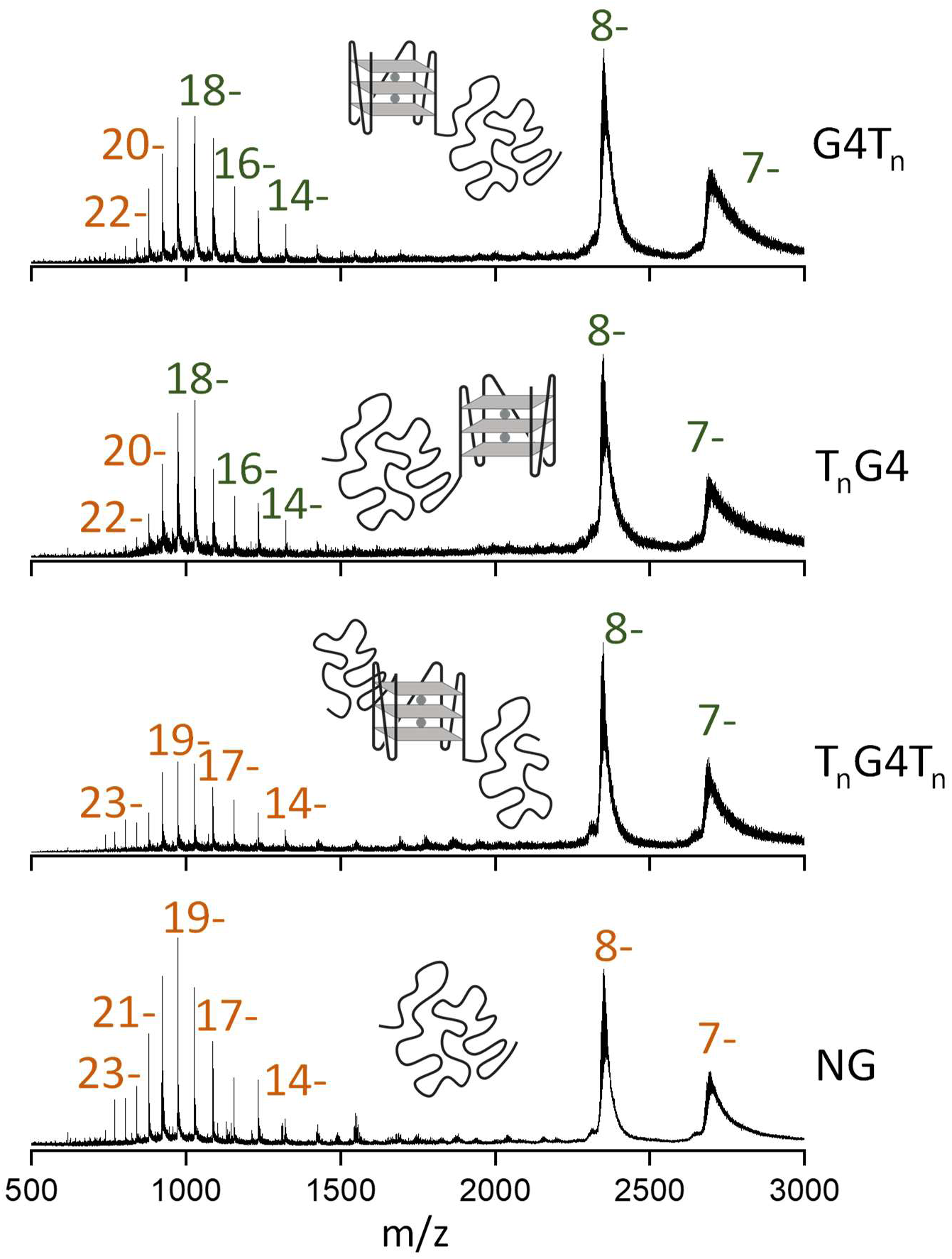
ESI-MS spectra of four 60-mer oligonucleotide structures: G4Tn, TnG4, TnG4Tn, and NG (top to bottom, 15 μM each) in 150 mM aqueous NH_4_OAc. MS data are filtered in the IM-MS browser to exclude background signals with low charges (1- to 4-). The unfiltered data are shown in Figure S2. Zooms on the adducts distributions are shown in Supporting Information Figures S3—S7.

Throughout all experiments, the fragmentor voltage was kept at 320 V, a value that allows to preserve the folded ammonium-bound structures of even very fragile G4s.^26^ In more detail, we distinguish: i) low charge states (7- and 8-), for which we distinguish many more adducts than the specific two expected (increasing the fragmentor voltages to 400 V or 450 V could fully desolvate ammonium adducts on the 8- ions but also led to the loss of internally bound cations, see supporting Figures S8 and S9); ii) some missing charge states (9- to 12-); iii) high charge states that still contain specifically 2 NH_4_^+^ ions (attributed to location at the specific coordination sites of the folded G4 subunits^38, 39^), centered on 16-for TnG4 and G4Tn; and iv) high charge states that do not contain NH_4_^+^ ions, centered on 18- for G4Tn and TnG4 and 19- for TnG4Tn and NG. TnG4Tn and NG display even higher charge states. This effect is much more pronounced at 50 mM aqueous NH_4_OAc (Figure S10). The extent of high charges thus depends on the presence of a quadruplex subunit (NG forms more of the high charge states, see Figure 1 and Figure S10) but also on the tail location: when a polythymine overhang is present at both ends, the CSD is similar to the NG sequence. Lowering the ionic strength to 1 mM NH_4_OAc (Figure S11) drastically favors higher charges. Both TnG4Tn and NG populate the highest charge states.

### Native IM-MS suggests beads-on-a-string gas-phase structures for G-quadruplex containing sequences

Next, we examined the CCSD of each charge state, for structures with intact (2 NH_4_^+^ ions bound) or disrupted (no NH_4_^+^ ions bound) G4 subunits (see Figure 2 for G4Tn). The average CCS increases with the number of charges, but groups also emerge. For example, the CCSDs of G4Tn show three distinct groups at 150 mM (Figure 2A). The first group comprising low charge states (7- and 8-, retaining two NH_4_^+^ ions plus non-specific ones), has CCS values below 1500 Å². A second group of charge states (9- to 13-) is missing. The third group shows CCS values between 2500 Å² and 3200 Å², while the mass indicates that two NH_4_^+^ cations are retained. The fourth group, lacking internally bound NH_4_^+^, exhibits CCS values above 3200 Å². The relative abundance of these groups changes with lowering the ionic strength. The first group dominates at 150 mM NH_4_OAc, the third group dominates at 50 mM NH_4_OAc (Figure 2B), the fourth group dominates at 10 mM NH_4_OAc (Figure 2C), and fully takes over at 1 mM, along with a few higher charge states (Figure 2D).

**Figure 2.**
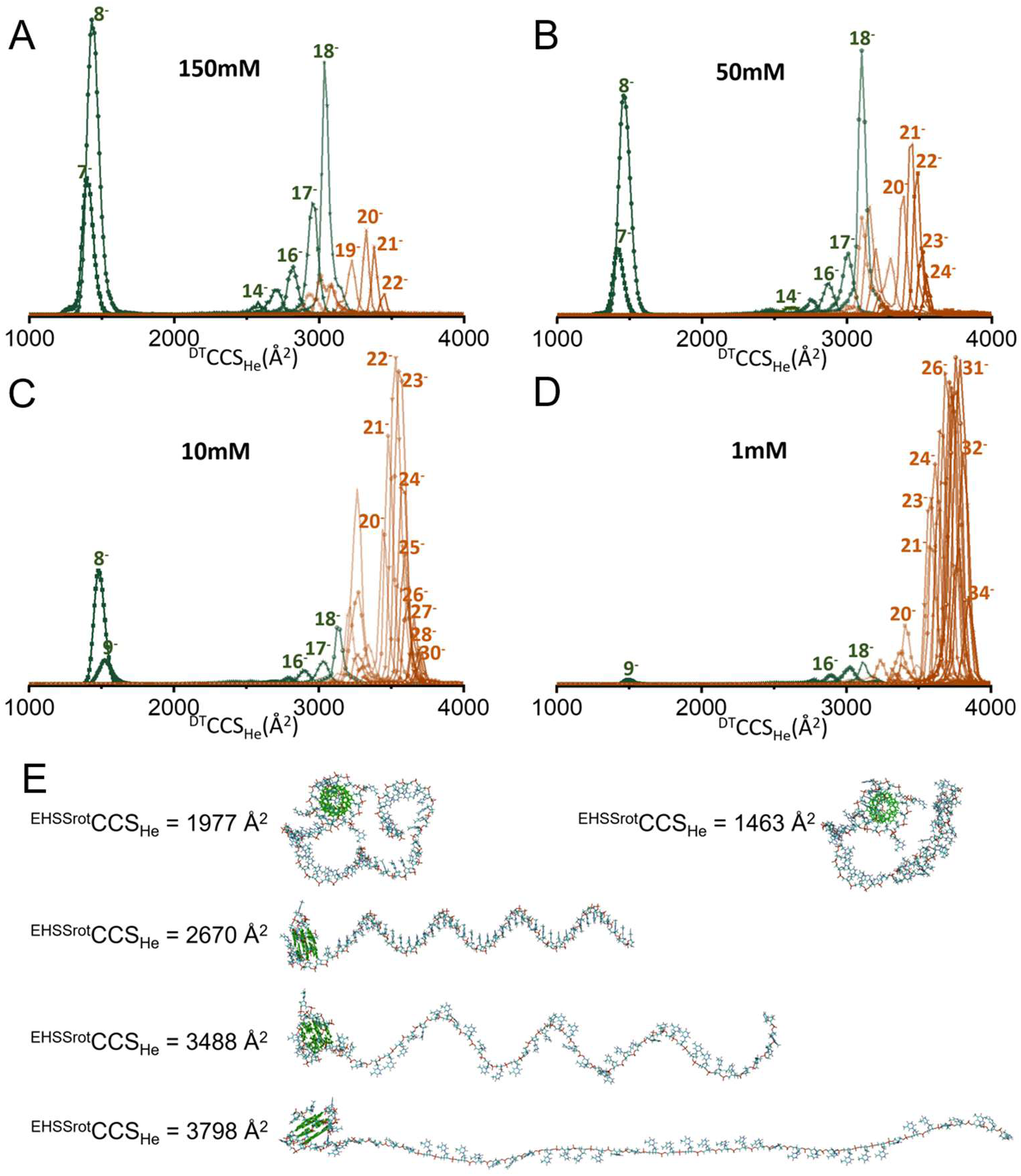
CCSD for the G4Tn oligonucleotide in various ionic strength, 150 (A), 50 (B), 10 (C), and 1 (D) mM aqueous NH_4_OAc. Ions with two specifically bound NH₄⁺ are in green, and ions without these bound cations are in brown. Some charge states appear both with and without two specific NH_4_^+^. Panel E shows five different conformation models of G4Tn in vacuo, all with a folded G4 subunit, along with their calculated CCS values, demonstrating that a wide range of CCS values are compatible with a preserved G4 subunit.

How can we imagine gas-phase conformations that are compatible with these observations? First, each set of 2 specifically bound NH_4_^+^ ions bound to the oligonucleotide, while the peaks corresponding to 1 or 3 NH_4_^+^ adducts are much lower, indicates that one G4 subunit is retained. However, the subunit region comprises 15 bases, while our strands are 60 bases long. The CCS indicates the global compactness of the gas-phase structures. By putting together these two pieces of information, we can infer the subunit folding status and the global unfolding (extension) status.

For reference, the Rayleigh limit radius of a water droplet that would have the same mass as that of the oligonucleotide (*ρ* = 1 g.cm^-3^) is *Z*_R_ = 10.8 to 10.9, and the corresponding fraction of charged phosphates (*Z*_R_/*P*) is thus 0.18. For 60-nucleotide sequence, CCS values below 1500 Å² obtained for charge states significantly below the Rayleigh limit (7-to 9-) should thus be formed via the CRM. Their CCS values corresponds to globular structures obtained by forming non-native hydrogen bonds between bases and phosphate groups (see Figure 51 in our review^40^). We thus postulate that the G4 subunit is folded and that the thymine tail wraps around it. Note that, given the oligonucleotide length and the unavailability of an appropriate gas-phase force field, it was difficult to generate satisfactory gas-phase structures by molecular modeling, and the models shown in Figure 2E serve more to visualize if experimental structures should be though of as more compact or more extended than some of the generated structures. For example, the structure having a CCS of 1977 Å² was generated by PM7 for a 8- ion, but was too large. The structure having a CCS < 1500 Å² was generated with the dT_24_ extremity constituted by four DFT models of dT_6_. Its CCS is closer to the experimental one, yet we don’t think it is realistic. Modeling progressive droplet desolvation (CRM) could bring us closer to realistic models, but this is out of the scope of the present work.

The next observed group has preserved G4 subunits (indicated by the 2 NH_4_^+^ ions), and the CCS values are compatible with relatively extended thymine overhangs. This kind of bead-and-string structure is compatible with the BEM ion production scenario. The next CCS distributions correspond to more extended structures with the G4 subunit unfolded. These fully extended structures could be formed either directly through a CEM ion production scenario, or initially via the BEM followed by Coulomb unfolding in the gas phase, leading to the loss of inner NH_4_^+^ cations and further extension. At physiological ionic strength, given that the folded fraction is close to 100% (Figure S1), we think the latter scenario is plausible, and CIU data on charge states 19- to 21- (see last section) show that the multimodal distributions comprise a low-CCS peak that is only present at low voltage. At higher charge states (CCS > 3500 Å²), the CEM scenario suffices to explain the observations.

TnG4Tn is an intriguing case: it does not display a high-charge state distribution with preserved ammonium adducts (Figure S12). Note, however, that experiments under native supercharging conditions (see last section) produced 14- and 15- charge states with preserved ammonium ions. The behavior without supercharging agents could be explained by two scenarios. (i) Ions are initially formed via the BEM in a similar fashion but then undergo gas phase unfolding and lose the ammonium ions. It is possible that a Coulomb-driven pulling effect is exerted on both ends of the G-quadruplex by the multiply charged polythymine tails, which may promote unfolding. (ii) Ion production must start from an overhang and thus by CEM, and once the process is started it keeps unfolding the central G4 during ion formation. Although the sequence TnG4Tn has the lowest solution thermal stability (Figure S1), the folded fraction predominates at room temperature. Molecular modeling (Figure S13) indicates that CCS values are compatible with an unfolded central G4 without the thymine chains fully extend. Our data does not allow us to choose one scenario over the other, but the ion production mechanism clearly depends on the positions of the folded subunit and unstructured overhangs.

Extended structures are prominent in the control NG, even at physiological ionic strength. CCSD produced by high charges indicates the presence of extended structures at 150 mM and 50 mM NH_4_OAc (Figure S14). At 1 mM aqueous NH_4_OAc, NG exhibits multimodal CCSD centered on 23-, 29-, and 32-, with CCS > 3500 Å² consistent with a fully extended conformation. This multimodal distribution might indicate that several ion production scenarios might still be at stake: CEM for the highest charge states and BEM (although the beads have no set localization) at slightly lower charge states.

In summary, IM-MS results confirm that 60-mers containing a single G-quadruplex can adopt conformations that contain both a folded G4 subunit and an extended polythymine overhang. The only ion production scenario compatible with this observation is the BEM. The CEM however becomes more prevalent at lower ionic strength, and for nonfolded structures. The position of the overhangs matters, however, as shown by the results obtained for TnG4Tn. We thus next examined a sequence with a G4 subunit at each end, and a polythymine spacer in the middle.

### Pre-folded beads at both termini: G4TnG4 shows unimodal CSD and CCSD in physiological ionic strength, and multimodal distributions at low ionic strength

Unlike single-G4 sequences, G4TnG4 forms only low charge states 7- and 8- ions at physiological ionic strength (Figure 3). Its unimodal CCSD (slightly below 1500 Å², Figure 4A) corresponds to a globular structure. The existence of two quadruplexes at the termini thus disfavors the production of high charge states with extension of central polythymine spacer. Even under native supercharging, only 7- to 9- charges were detected in 150 mM NH_4_OAc (Figure S17). Thus, at physiological ionic strength, the CRM ion production scenario predominates in all cases. Stacking between the two G4 subunits is also possible,^41,42^ keeping the overall fold compact in solution and even preventing supercharging.

**Figure 3.**
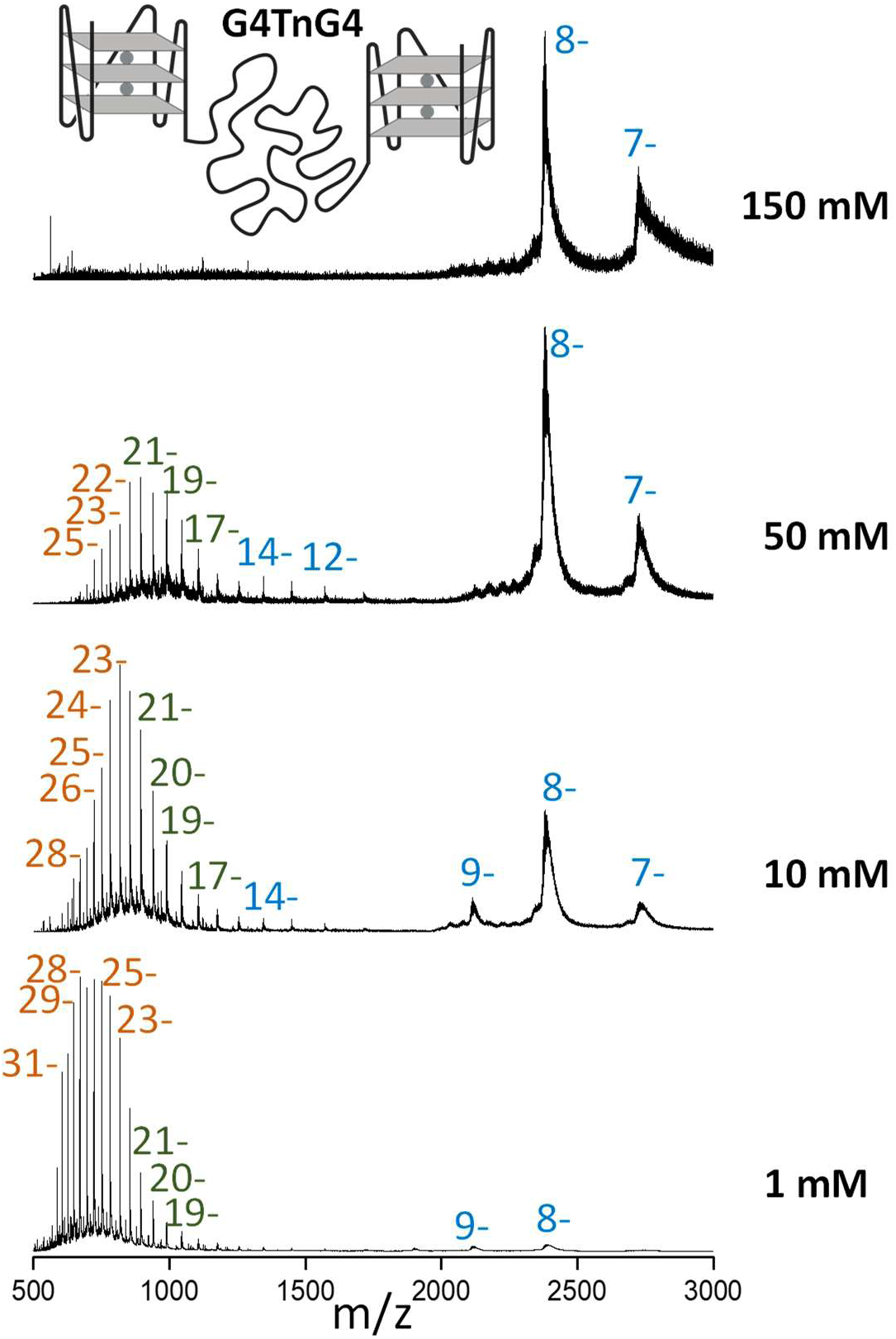
ESI-MS spectra of the G4TnG4 oligonucleotide, comprising one quadruplex subunit at each end, measured at four ionic strengths (150, 50, 10, and 1 mM, top to bottom, 15 μM each). Unfiltered spectra are shown in Figure S15. Zooms on the adducts distributions are shown in Supporting Information Figure S16.

**Figure 4.**
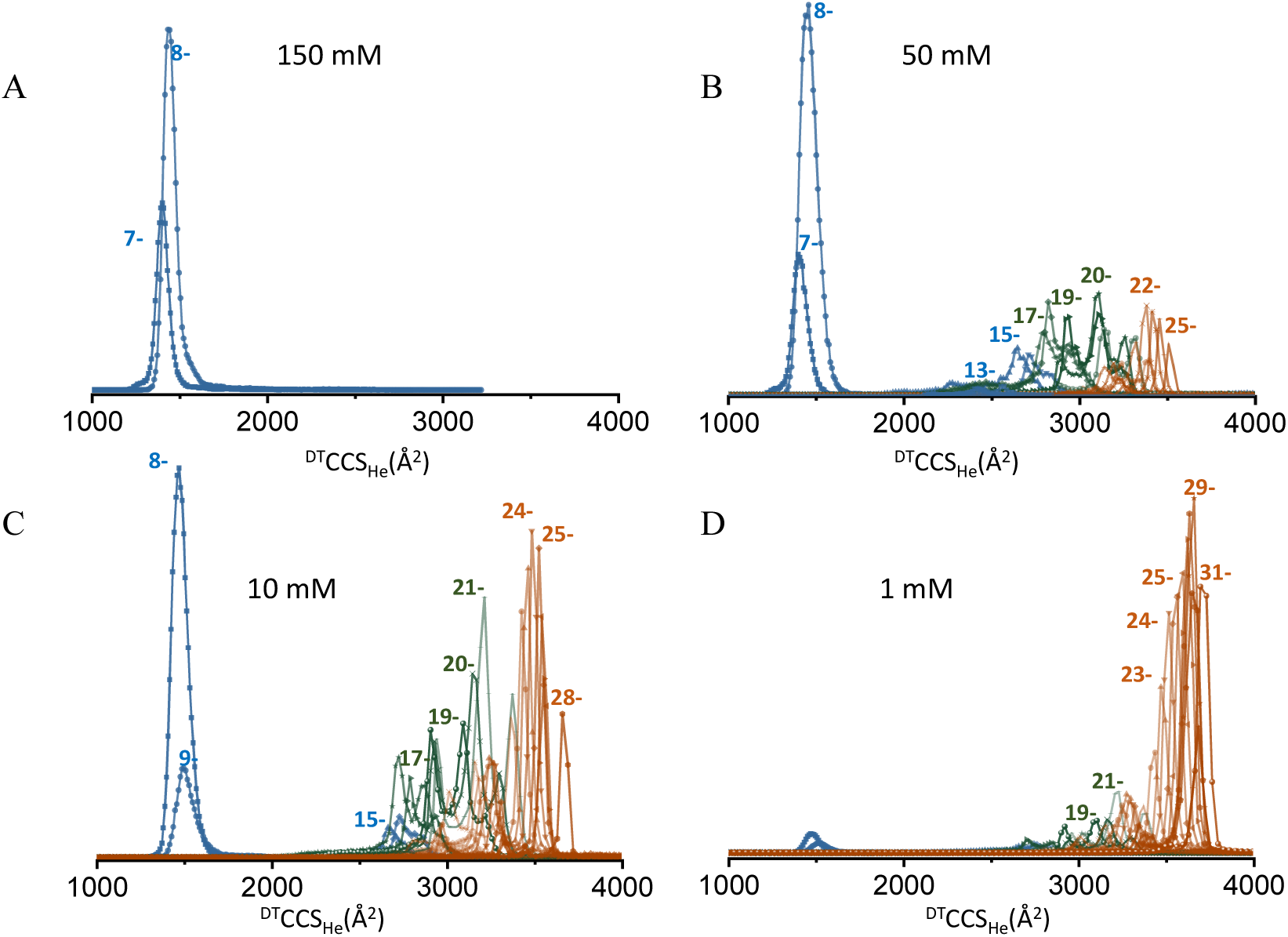
CCSD of G4TnG4 oligonucleotide in various ionic strengths, 150 (A), 50 (B), 10 (C), and 1 (D) mM aqueous NH_4_OAc. Ions with four specifically bound NH₄⁺ are in blue, two bound cations are in green, and ions without bound cations are in brown.

At 50 mM NH_4_OAc, we begin to observe ions with higher charge states (12- to 14-) which are partially extended, yet retain their 2 G4 subunits, as indicated by the preservation of mainly 4 NH_4_^+^ ions (see Figure S16). Charge states 15- and 16- undergo ammonia loss, two at a time. The CCSDs (blue in Figure 4) are between 2200 and 2800 Å². Again, the only scenario that can lead to such observation is the BEM. The next distributions include structures that retain only 2 NH_4_^+^ ions and therefore one G4 subunit (green), then those that have lost both G4s and are the most extended (brown). Only the latter observations are compatible with the CEM. Again, the fraction of the analyte population that is ionized through via a CRM, BEM or CEM pathway depends on the ionic strength.

### The competition between CRM, BEM and CEM ion production scenarios depends on the position of subunits and unstructured overhangs

Up to now, ion formation scenarios of oligonucleotide have been described as a competition between CRM and CEM.^10^ The present study with longer sequences containing folded domains and unstructured linkers suggests that the BEM process, which is hybrid between CRM and CEM, can also happen. Figure 5 conceptualizes this continuum.

**Figure 5.**
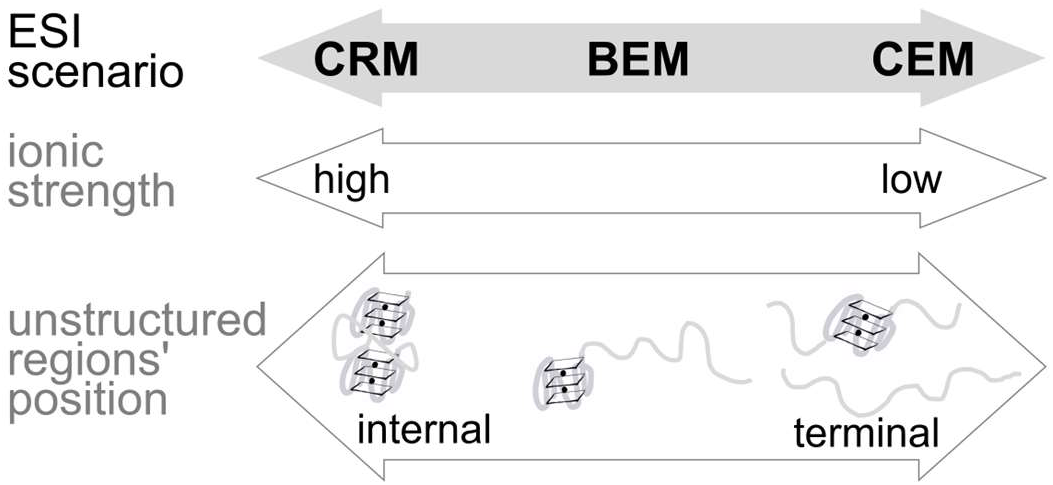
Ionic strength and position of unstructured regions are the major factors influencing the nucleic acids ion production mechanism in negative ion mode. The bead ejection mechanism (BEM) is intermediate and hybrid between the charged residue mechanism (CRM) leading to globular structures and the chain ejection mechanism (CEM) leading to fully extended structures.

High ionic strength favors the CRM, while low ionic strength favors the CEM. Unstructured terminal regions favor the CEM, while folded domains on the termini favor the CRM. Folded G-quadruplex domains at both termini favor the CRM, which becomes the only ion production scenario at physiological (high) ionic strength. However, charging beyond the Rayleigh limit becomes possible by lowering the ionic strength, extension becomes inevitable and beads-on-a-string structures appear. Without supercharging agents, CRM at these charge states is not realistic, and thus ions can only be produced by the BEM. When one terminus consists of a G-quadruplex domain and the other by an unstructured region, the BEM is observed even at physiological ionic strength.

Surprisingly, for TnG4Tn, the G-quadruplex domain remains folded only in the low charge states produced by the CRM. None of the peaks at charge states 13- or higher contained inner ammonium ions indicative of a preserved G-quadruplex. Also, the charge states were on average higher with this sequence: CSD and CCSDs were similar to the unstructured control. We reason that the CEM ionization pathway must start from one of the ends, and that once the CEM process has started at an extremity, it could induce the unfolding during ionization. Alternatively, the BEM could be at stake, but due to a Coulomb pulling effect from the poly(T) overhangs attached at each extremity of the G4 subunit and charged by CEM, the subunit could be disrupted.

### Charge-to-phosphate ratios between 0.25 and 0.33 is optimal to discriminate folded domains preserved from solution

Another interesting question is: what charge states are the most useful to obtain information on the solution structures? As reported before, the lowest charge states produced by the CRM are not particularly informative because the ion structures are equally compact and globular. In our 60-mers, these charge states correspond to a charge density (charge-to-phosphate ratio, *z*/*P*) of 0.11 to 0.14. Recall that the Rayleigh limit charge density in water is around 0.18. The highest charge states (21- and more, i.e. *z*/*P* > 0.35), are equally uninformative, this time because the G4 subunits are disrupted in the gas phase, as indicated by the loss of inner NH_4_^+^ cations.

We therefore attempted to promote intermediate charge states using supercharging agents that favor CRM while imparting more charges,^43^ making the ions more suitable for collision-induced unfolding experiments.^14^ For all 60-mers, charge states 9- to 13- are notably absent under native conditions. Here, adding 0.4% PC likewise reveals otherwise hidden charges from 9- to 12- for instance for G4Tn (Figure S18), and their CCS increase gradually with the charge state. Furthermore, for TnG4Tn, a low-abundance population of 13- to 15- ions retaining two bound NH_4_^+^ ions emerges under native supercharging conditions, as shown in Figure S19. This supports Coulomb-induced unfolding of the overhangs away from the G4 core. This also suggests that the high-charge state distribution observed in native conditions without supercharging agents were produced via a different mechanism. Only the sequence G4TnG4 could not be supercharged from physiological ionic strength (Figure S17), so we could only use high charge state data obtained at lower ionic strength.

We found that the most discriminative region for our 60-mers consists of charge states 15-to 20- (0.25 < *z*/*P* < 0.35). Figure 6 shows how the collision cross section of the peak maxima in the softest conditions (all CIU data are shown in supporting information Figures S20—S25) depends on the sequence at those charge states. Supplementary Figure S26 shows an analogue of Figure 6, using data obtained in presence of supercharging agent, leading to similar conclusions. The structures containing two preserved G-quadruplexes are significantly more compact than those that contain only one, and those that contain none (NG control). This discrimination based on CCS is made possible by the BEM ionization scenario. Interestingly, when G4TnG4 contains only two ammonium ions and not four at charge states 17- to 20-, it is more compact than G4Tn or TnG4 with two ammoniums, so despite the partial loss of ammonium ions there is a memory of the presence of two folded subunits.

**Figure 6.**
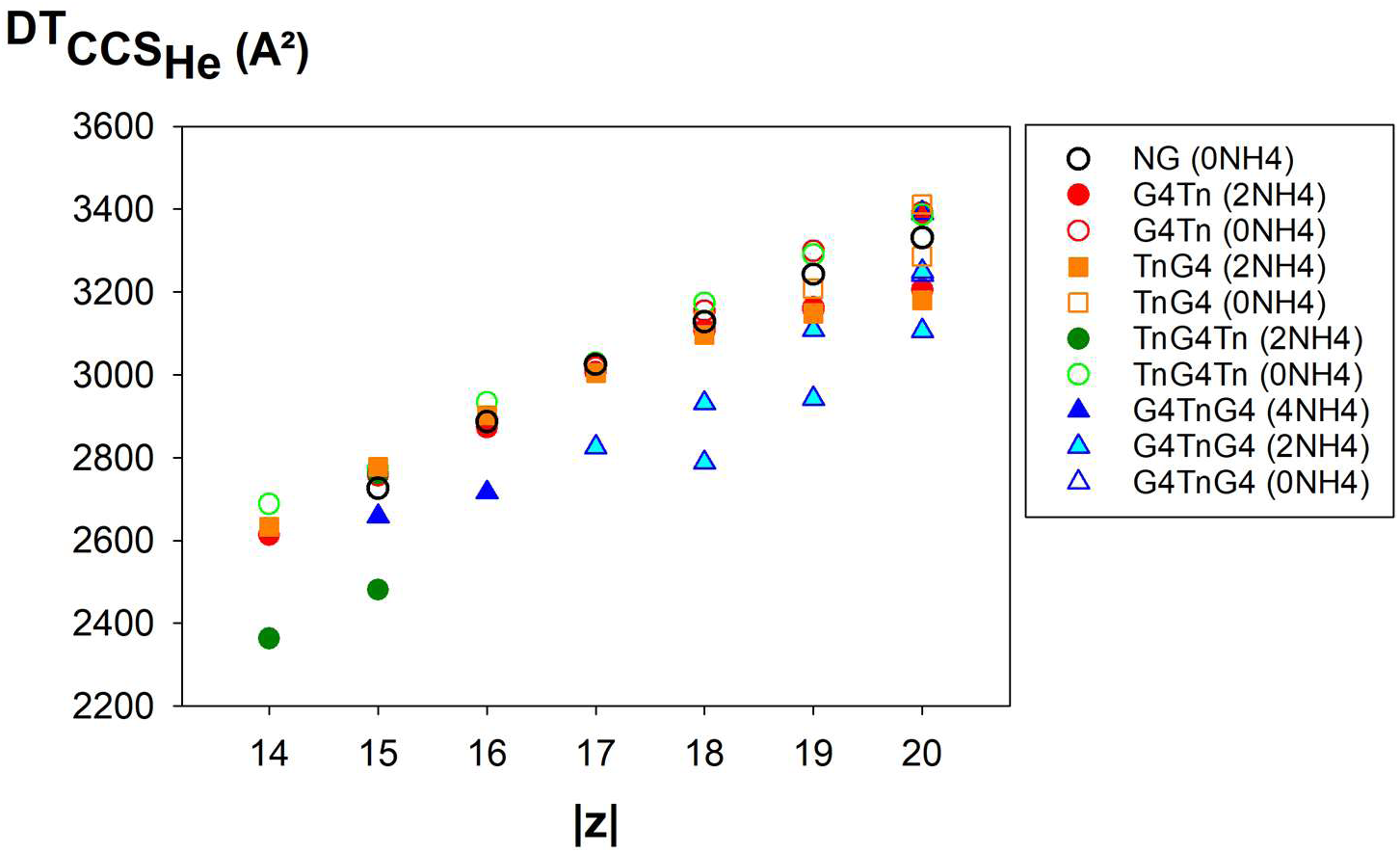
Helium collision cross sections of the 60-mer DNAs, recorded from aqueous 50 mM NH_4_OAc, except for TnG4Tn with 2 NH_4_^+^ 14- and 15-, which was recorded in 150 mM NH_4_OAc and 0.4% propylene carbonate. Filled symbols correspond to sequences with all ammonium ions and thus with preserved G-quadruplex structures.

However, those charge states are not always present when infusing the samples from purely aqueous NH_4_OAc solutions at physiological ionic strength. Supercharging agents do not always succeed at charging to the optimum level. It can be useful in specific cases, such as TnG4Tn here, to produce some charge states with the CRM and have a chance to observe folded domains in low-energy conditions. In contrast, it is always possible to lower the ionic strength in aqueous solution to generate the most informative charge states for ion mobility analysis, while favoring the BEM.

## Conclusions

We presented here several gas-phase structures (elongated but with inner ammoniums indicating preserved G4 folded domains) which can only have a bead on a string structure. Our study confirms that the bead ejection mechanism (BEM), which should be seen as a charged residue scenario for the folded regions and a chain ejection scenario for some of the unstructured regions, can also apply to nucleic acids. Unstructured regions located at the 5’ or the 3’-end favor the CEM, whereas the CRM is favored when the strand has folded domains on both termini. In future work, it would be interesting to investigate whether the sequence of the unstructured regions plays a role, and if the model applies to other kinds of folds than G-quadruplexes.

Favoring the BEM scenario is particularly interesting for native MS studies of nucleic acid folding in solution, because it leads to higher levels of charging. We found that a charge-to-phosphate ratio 0.25 < *z*/*P* < 0.35 is optimal to distinguish the solution folds based on the collision cross section. The CRM produces too low charge states, wherein folded domains and unstructured regions rather form extra non-native hydrogen bonds with one another,^21^ resulting in indistinguishably compact structures. This contrasts with the usual tenet according to which the lowest charge states are always the best for native MS.

Charge state tuning can be achieved either through the addition of supercharging agents or through lowering the ionic strength. The latter approach worked better. Adding supercharging agents did not always succeed at producing higher charge states, and the gradual charge state increase observed is more in line with the CRM. However, the results on TnG4Tn, where the G-quadruplex domain is flanked by two unstructured thymine overhangs, show that the BEM sometimes fails at preserving the folded domains. Therefore, results should be interpreted with caution, and incorporating well-designed control sequences remains essential for reliable conclusions.

## Acknowledgements

This work was funded by the Agence National de la Recherche (project ANR-18-CE29-0013 POLYnESI to VG and FR).

## Supplemental data description

Solution UV-melting curves, unfiltered MS data, zooms of the MS spectra showing the ammonium adducts distributions, supplementary MS spectra at other ionic strengths, supplementary collision cross section distributions, molecular model of unstructured TnG4Tn, ESI-MS data obtained under native supercharging conditions, CIU experiments, i.e., CCSDs as a function of the pre-IMS fragmentor voltage, equivalent of Figure 6 recorded in presence of supercharging agent instead of decreasing the ionic strenght (PDF).

