## supporting information for "Bead-Ejection Scenario in Electrospray Ionization of Multi-Domain Nucleic Acids"

### Supporting Information Table of Contents

| Number | Description | Page |
| --- | --- | --- |
| **Figure S1** | Melting curve of different 60-mers | 3 |
| **Figure S2** | ESI-MS spectra of G4Tn and TnG4Tn in high ionic strength (unfiltered data) | 4 |
| **Figure S3** | MS spectra of 7-, 8- ions of all oligonucleotides showing adducts | 5 |
| **Figure S4** | MS spectra of all high charge states of G4Tn showing adducts | 6 |
| **Figure S5** | MS spectra of all high charge states of TnG4 showing adducts | 7 |
| **Figure S6** | MS spectra of all high charge states of TnG4Tn showing adducts | 8 |
| **Figure S7** | MS spectra of all high charge states of NG showing adducts | 9 |
| **Figure S8** | MS spectra of 8- ion of G4Tn at F400 and F450 | 10 |
| **Figure S9** | Arrival time distribution of G4Tn at 8- charge state | 11 |
| **Figure S10** | ESI-MS spectra of 60-mers at 50 mM ionic strength | 12 |
| **Figure S11** | ESI-MS spectra of 60-mers at 10- and 1-mM ionic strength | 13 |
| **Figure S12** | CCSD of TnG4Tn at various ionic strengths | 14 |

| | | |
|---|---|---|
| **Figure S13** | Modelled structure of TnG4Tn | 15 |
| **Figure S14** | CCSD of NG at various ionic strengths | 16 |
| **Figure S15** | ESI-MS spectra of G4TnG4 at 150 and 50 mM ionic strength (unfiltered data). | 17 |
| **Figure S16** | MS spectra of all charge states of G4TnG4 showing adducts | 18 |
| **Figure S17** | Native supercharging of G4TnG4 | 19 |
| **Figure S18** | Native supercharging of G4Tn | 20 |
| **Figure S19** | Native supercharging of TnG4Tn and NG | 21 |
| **Figure S20** | CIU experiments for G4Tn and TnG4Tn | 22 |
| **Figure S21** | CIU experiments for G4Tn | 23 |
| **Figure S22** | CIU experiments for TnG4 | 24 |
| **Figure S23** | CIU experiments for TnG4Tn | 25 |
| **Figure S24** | CIU experiments for NG | 26 |
| **Figure S25** | CIU experiments for G4TnG4 | 27 |
| **Figure S26** | CCS of detected peaks as a function of the charge states, recorded in native supercharging conditions | 28 |

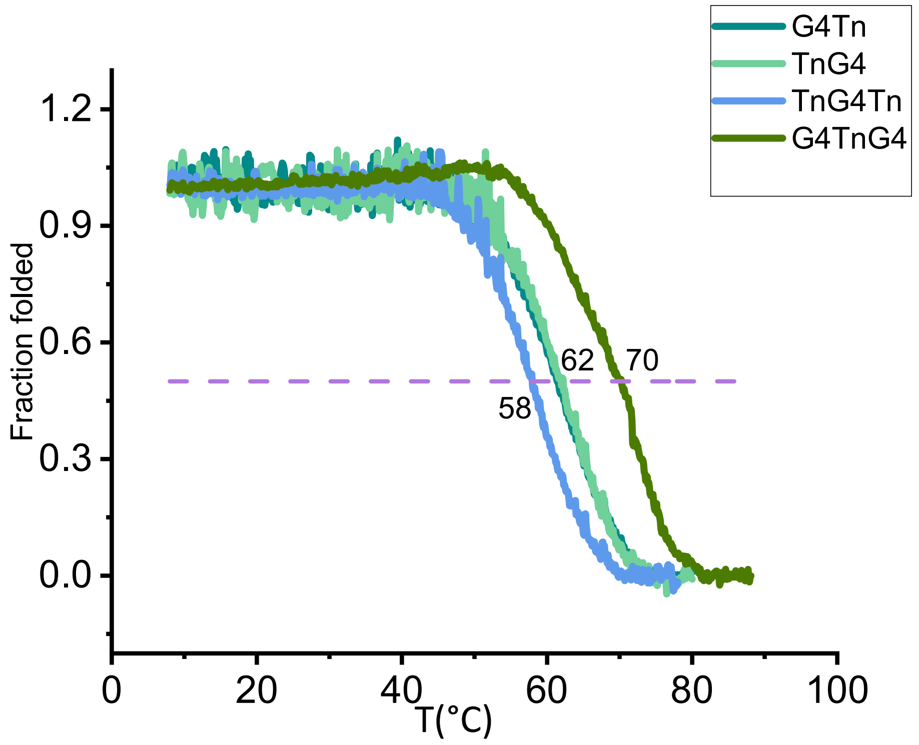


**Figure S1:** Melting curve of different 60-mer oligonucleotides at physiological ionic strength (150 mM aqueous $NH_4OAc$ solution) showing that the structures are folded at room temperature.

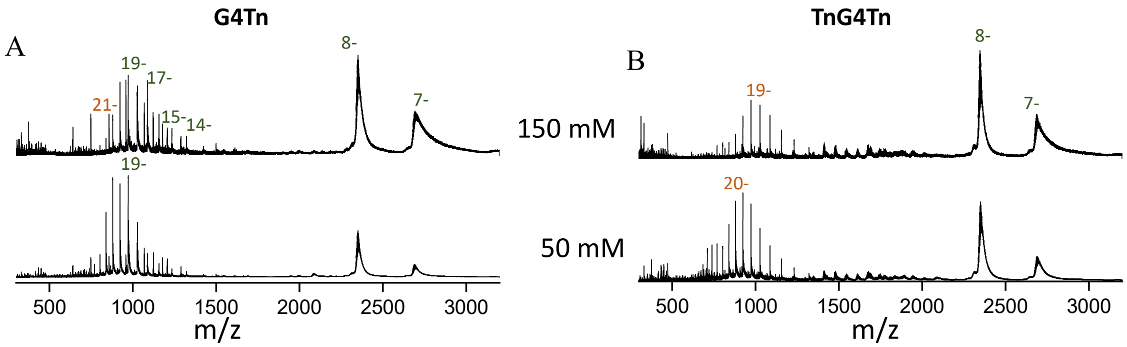


**Figure S2:** ESI-MS spectra of 60-mer oligonucleotides, G4Tn (A) and TnG4Tn (B) in 150 and 50 mM aqueous $NH_4OAc$ (unfiltered data).

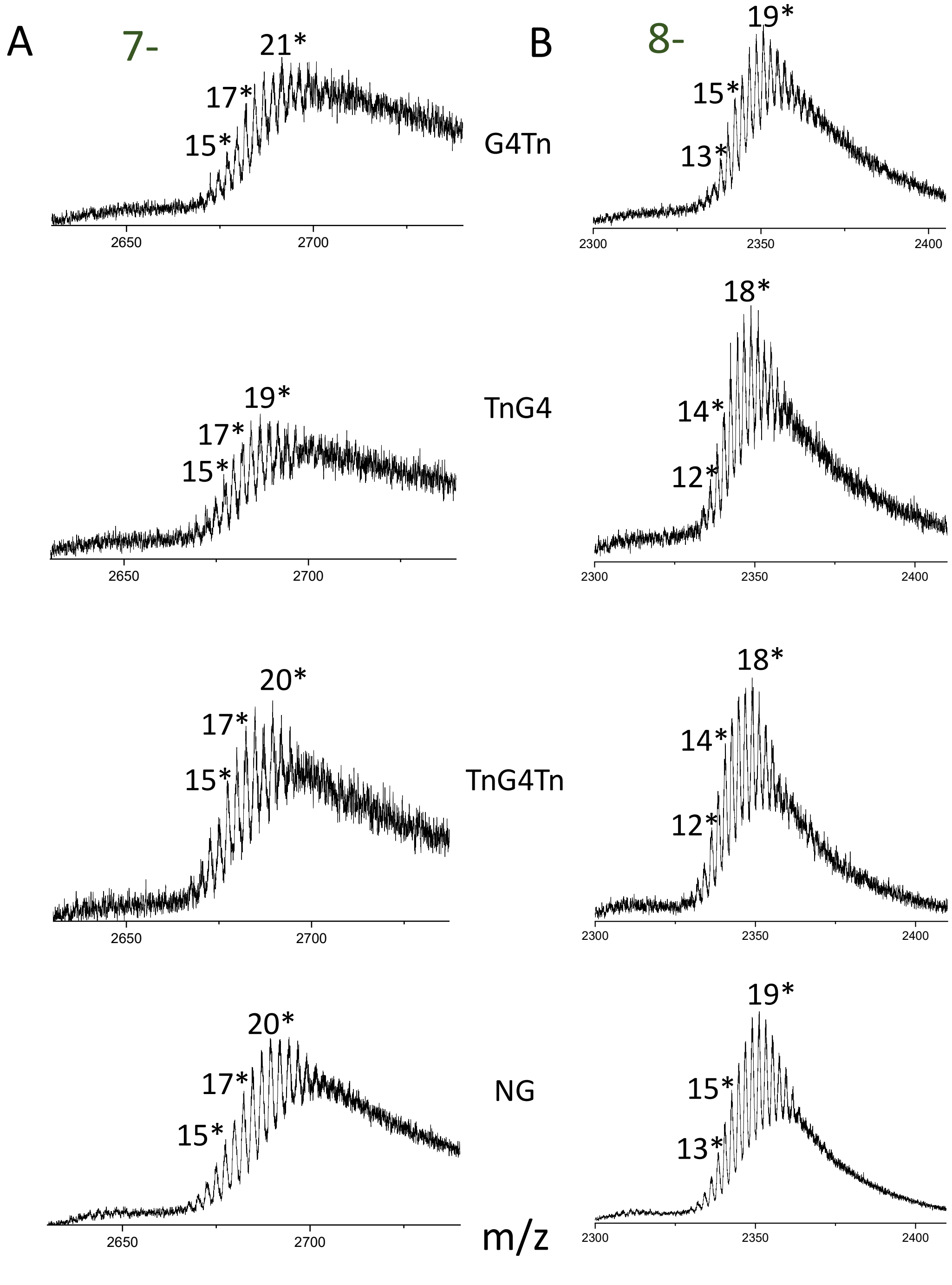


**Figure S3:** MS spectra of G4Tn, TnG4, TnG4Tn, and NG recorded at 320V exhibit multiple $NH_4^+$-adducts in the $7^-$ (A) and $8^-$ (B) charge states in 150 mM $NH_4OAc$. The numbers indicate the count of non-specific $NH_4^+$-adducts (marked with an asterisk *).

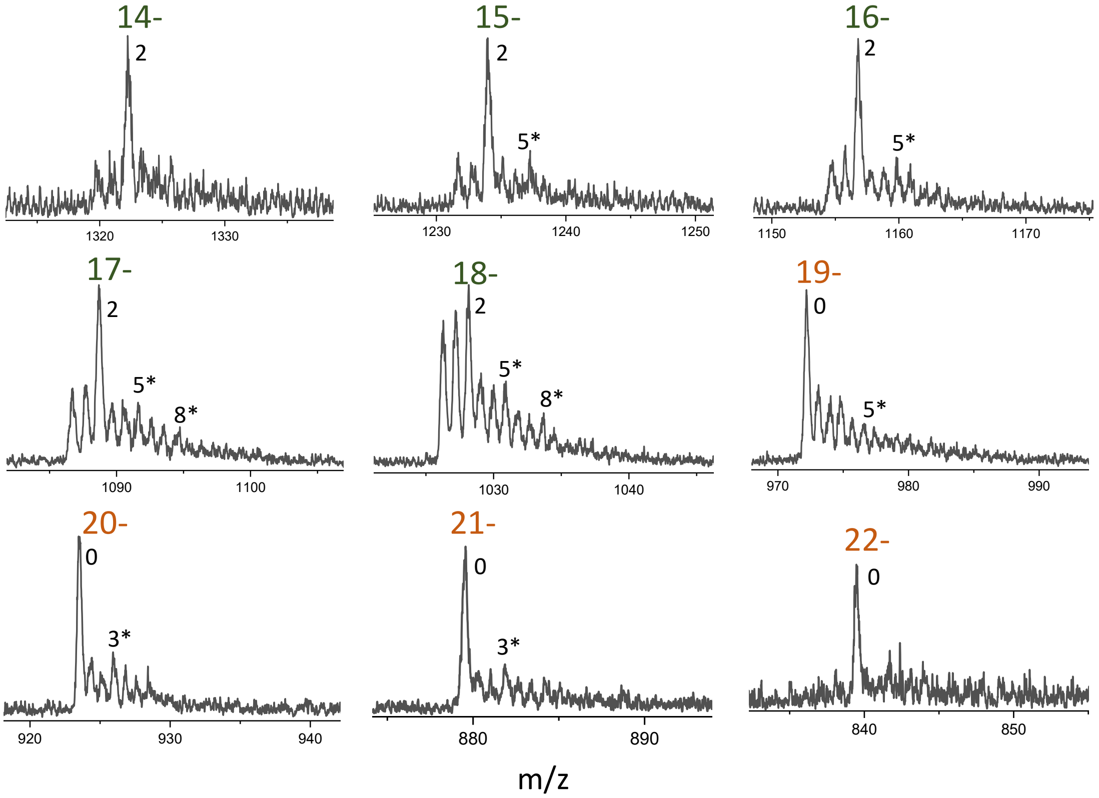


**Figure S4:** MS of G4Tn recorded at 320V show few $NH_4^+$-adducts at high charge states in 150 mM $NH_4OAc$. The numbers indicate the count of non-specific $NH_4^+$-adducts (marked with an asterisk *), and specific adducts (without asterisk).

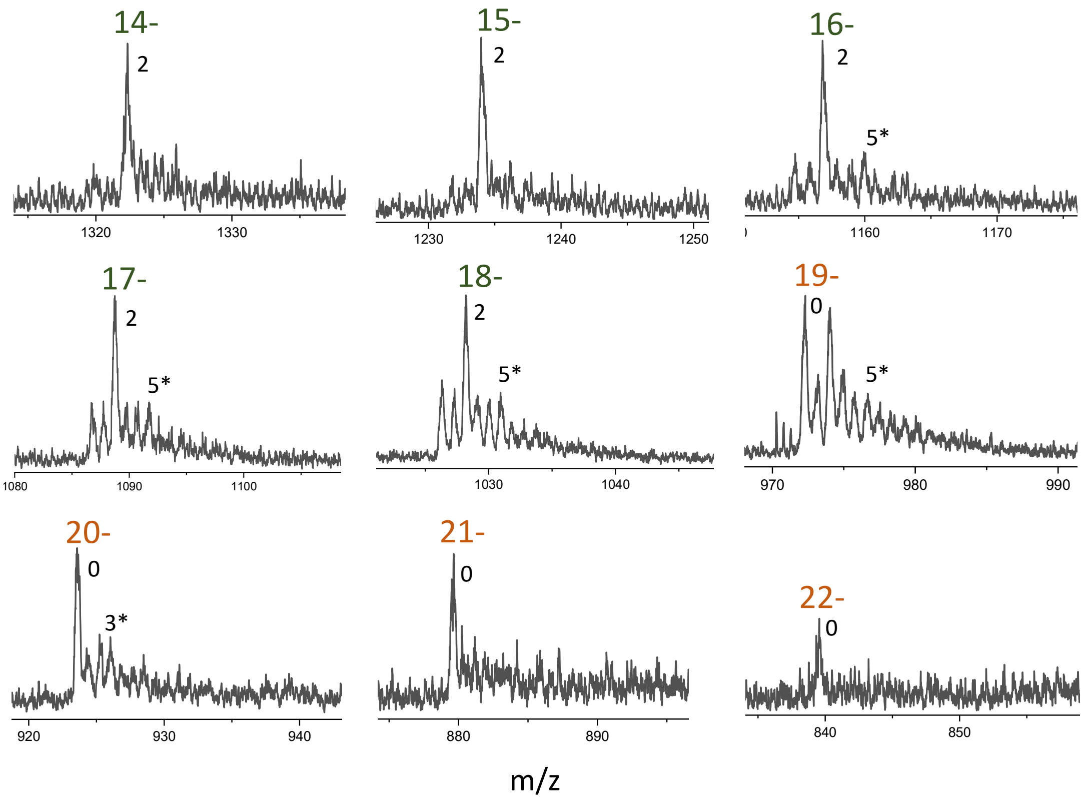


**Figure S5:** MS of TnG4 recorded at 320V show few $NH_4^+$-adducts at high charge states in 150 mM $NH_4OAc$. The numbers indicate the count of non-specific $NH_4^+$-adducts (marked with an asterisk *), and specific adducts (without asterisk).

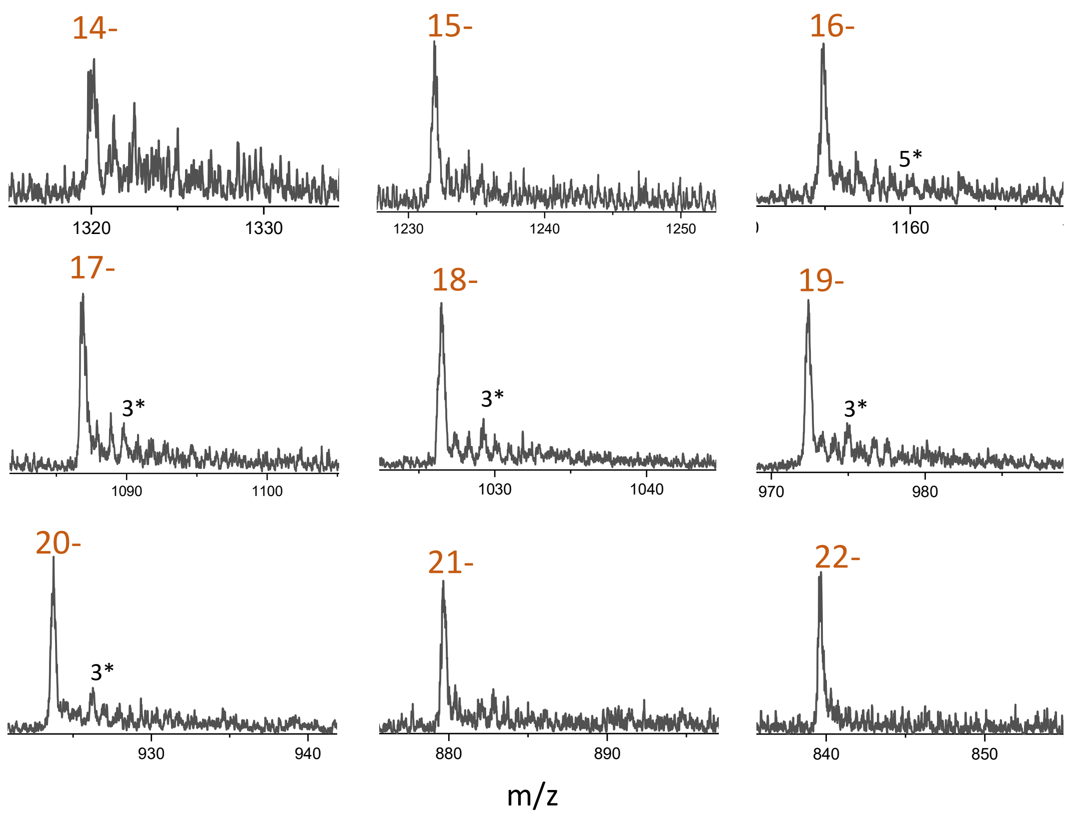


**Figure S6:** MS of TnG4Tn recorded at 320V show few $NH_4^+$-adducts at high charge states in 150 mM $NH_4OAc$. The numbers indicate the count of non-specific $NH_4^+$-adducts (marked with an asterisk *).

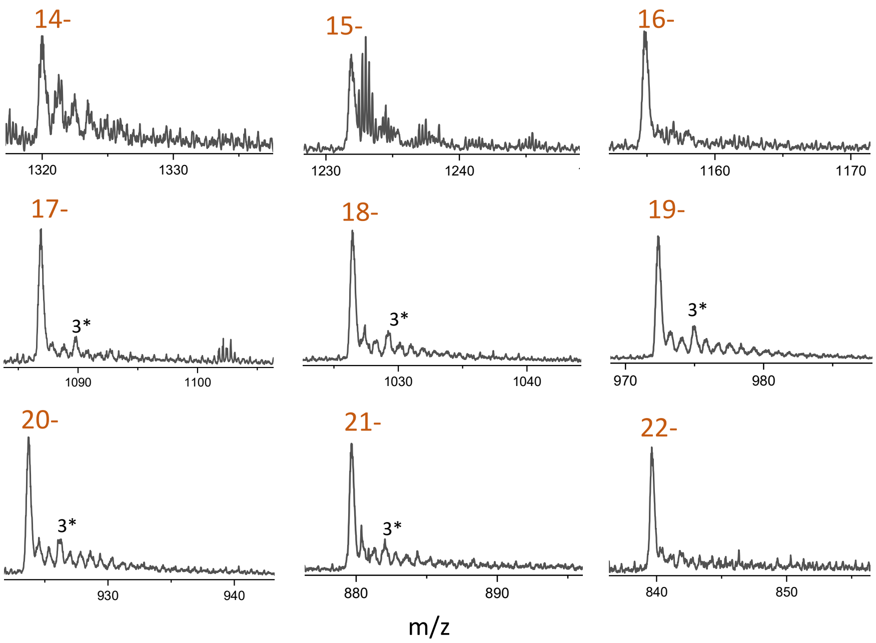


**Figure S7:** MS of NG recorded at 320V show few $NH_4^+$-adducts at high charge states in 150 mM $NH_4OAc$. The numbers indicate the count of non-specific $NH_4^+$-adducts (marked with an asterisk *).

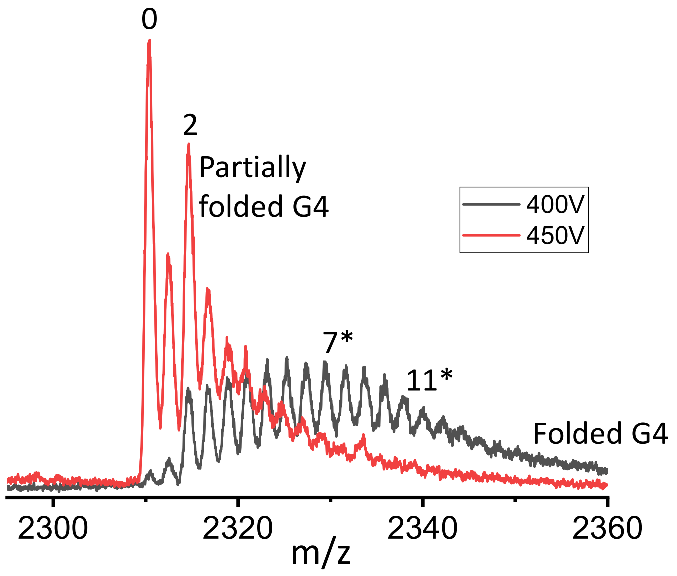


**Figure S8:** MS of G4Tn in 150 mM aqueous $NH_4OAc$ show multiple $NH_4^+$-adducts at a low fragmentor voltage (400V) in the low charge state, 8-. The numbers indicate the count of specifically bound $NH_4^+$ ions, while number marked with an asterisk (*) denote non-specific $NH_4^+$-adducts. Increasing the fragmentor voltage to 450V improves the removal of nonspecific adducts but also causes the loss of intact G4 structures, even at the low-charge state (8−).

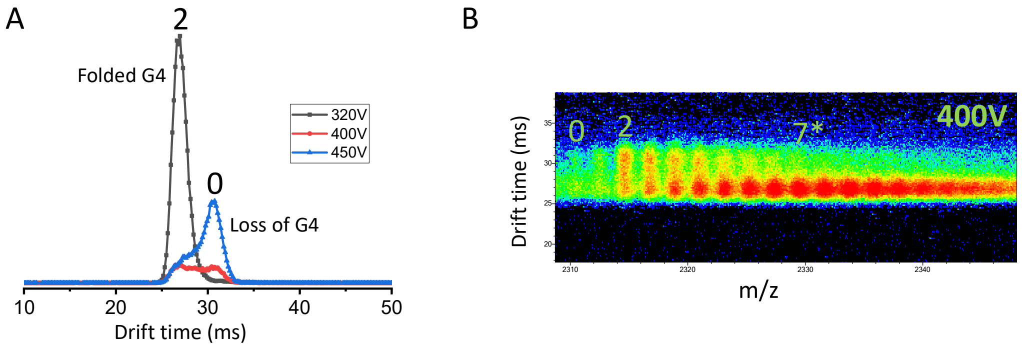


**Figure S9:** A) The arrival time distribution plot for G4Tn at an 8- charge state in 150 mM aqueous $NH_4OAc$ illustrates the effect of varying the fragmentor voltages. At 320V (low voltage), the structure is fully folded, appearing as a single distribution. At 400V, two distributions emerge: one with two specifically bound $NH_4^+$ ions and another lacking $NH_4^+$ ions. At 450V (high voltage), the distribution without bound $NH_4^+$ ions dominates, indicating the loss of G-quartets. B) The 2D IMS plot shows multiple $NH_4$-adducts at a fragmentor voltage (400V) in the 8- charge state. The numbers indicate the count of specifically bound $NH_4^+$ ions, while number marked with an asterisk (*) denote non-specific $NH_4^+$-adducts.

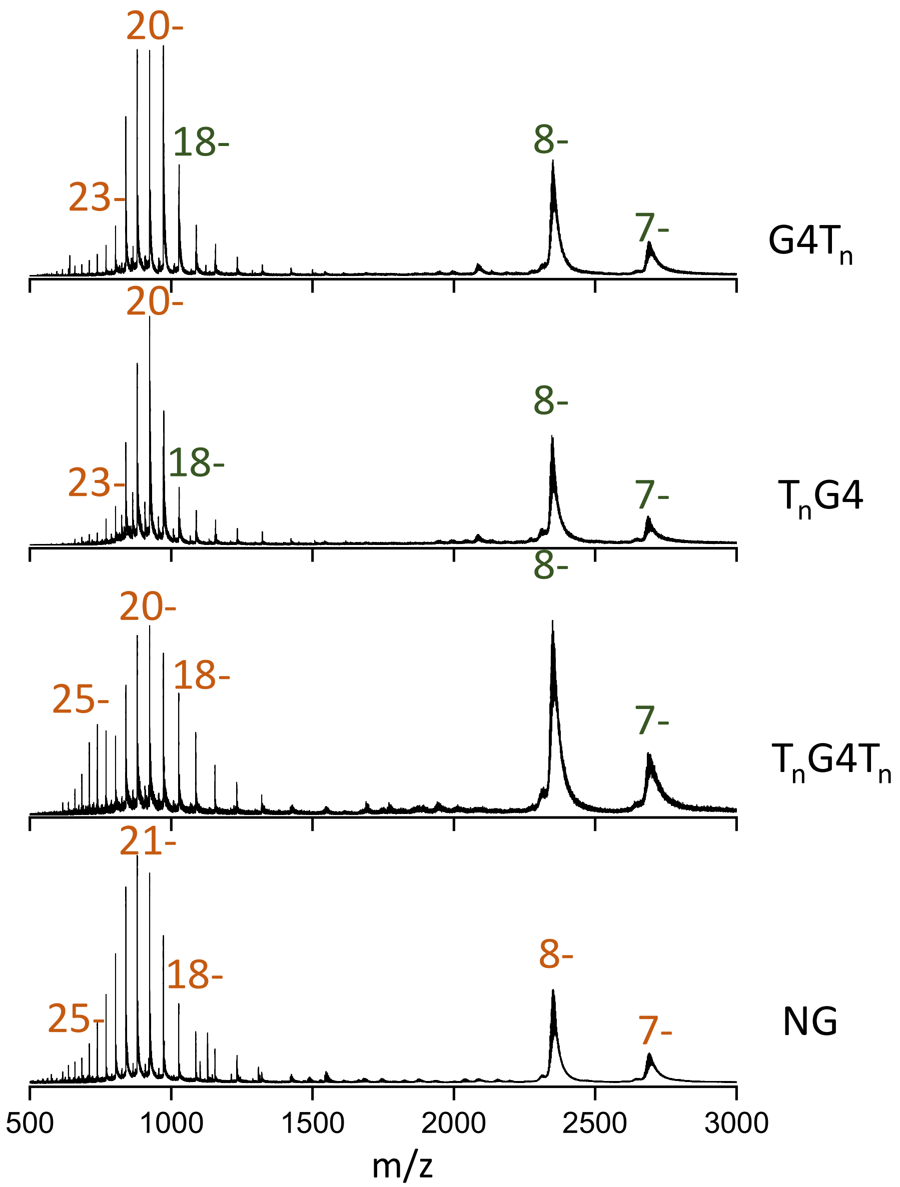


**Figure S10:** ESI-MS spectra of different 60-mer oligonucleotides (15 μM each) in 50 mM aqueous $NH_4OAc$.

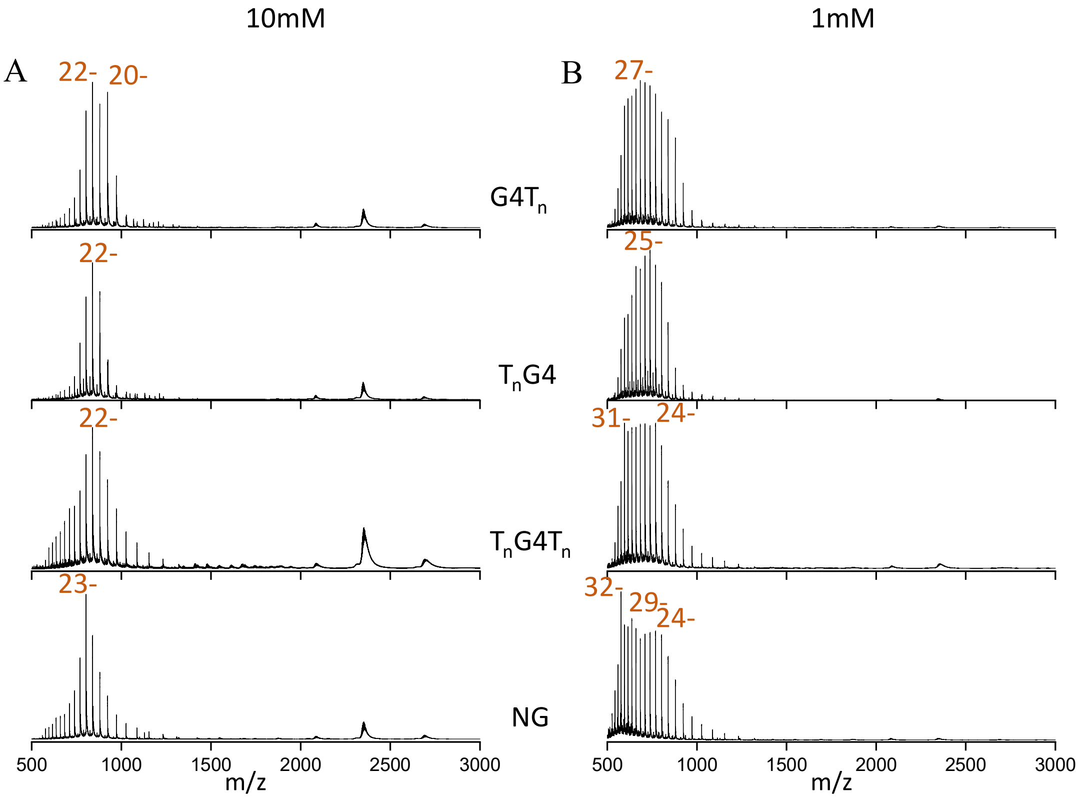


**Figure S11:** ESI-MS spectra of different 60-mer oligonucleotide structures (15 µM each) at low ionic strength, i.e., 10 (A), and 1 mM (B) aqueous $NH_4OAc$.

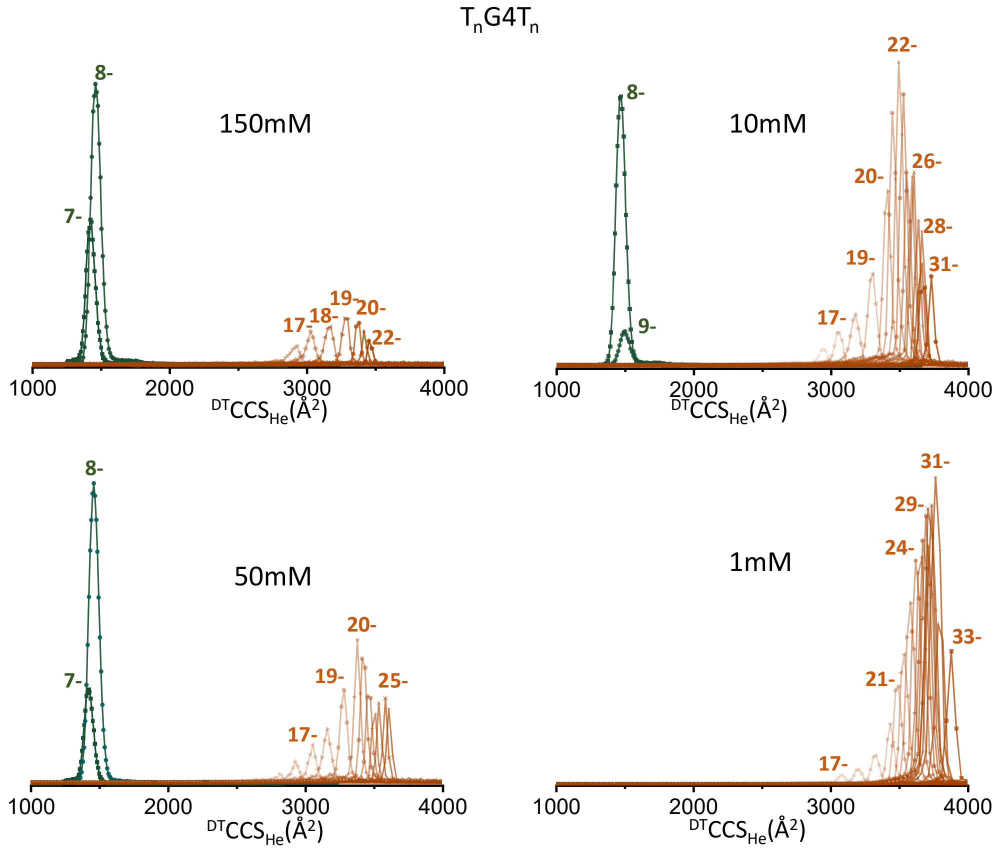


**Figure S12:** CCSD for ions of TnG4Tn at various ionic strengths showing the absence of third distributions of CCS.

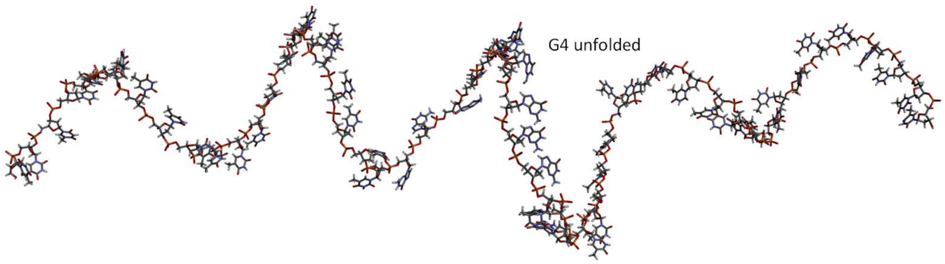


$^{Theo}CCS_{EHSSrot}$ = 3862.7 Å$^2$

**Figure S13:** Modelled structure of $(TnG4Tn)^{29-}$ shows the disruption of G4 core before complete elongation of thymine chains.

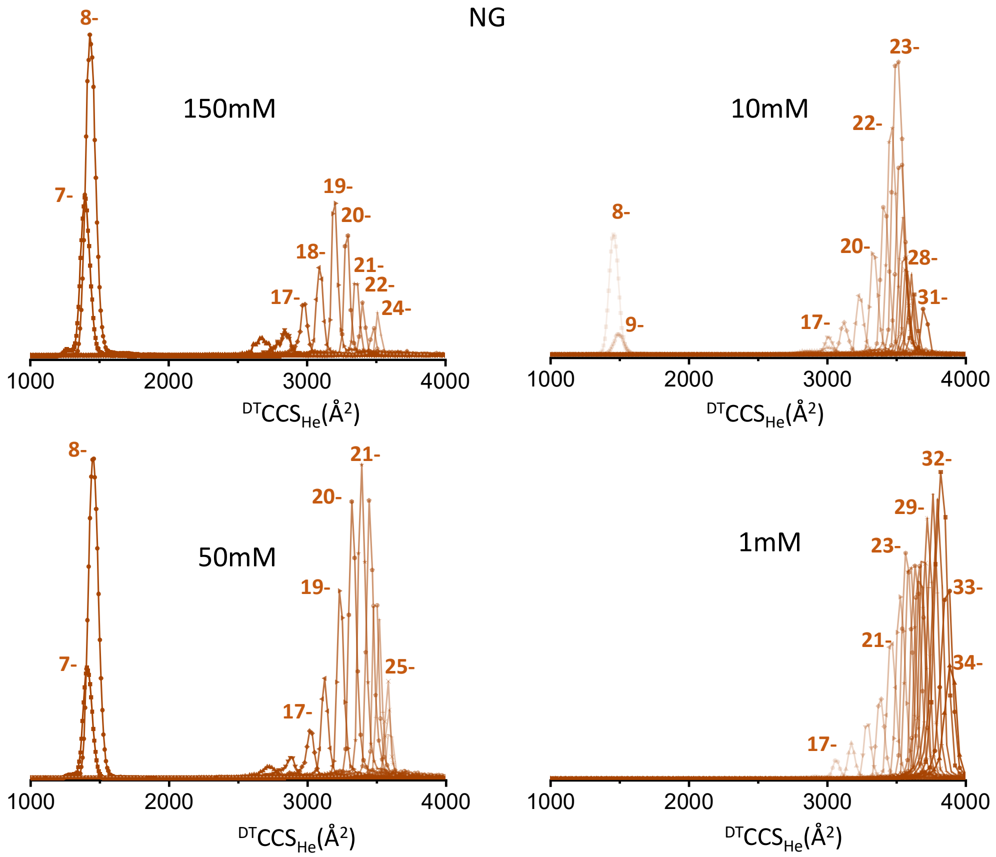


**Figure S14:** CCSD for ions of NG at various ionic strengths. The relative abundance of extended structures is higher (even at 50 mM ionic strength) in comparison to other 60-mers.

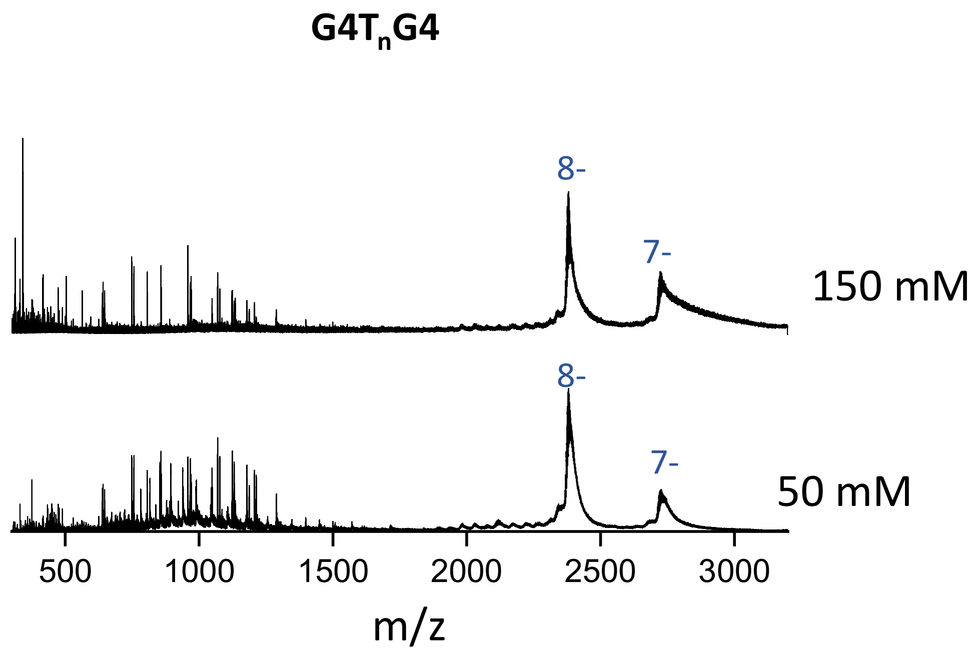


**Figure S15:** ESI-MS spectra of G4TnG4 at 150 mM and 50 mM aqueous $NH_4OAc$ solution (unfiltered data).

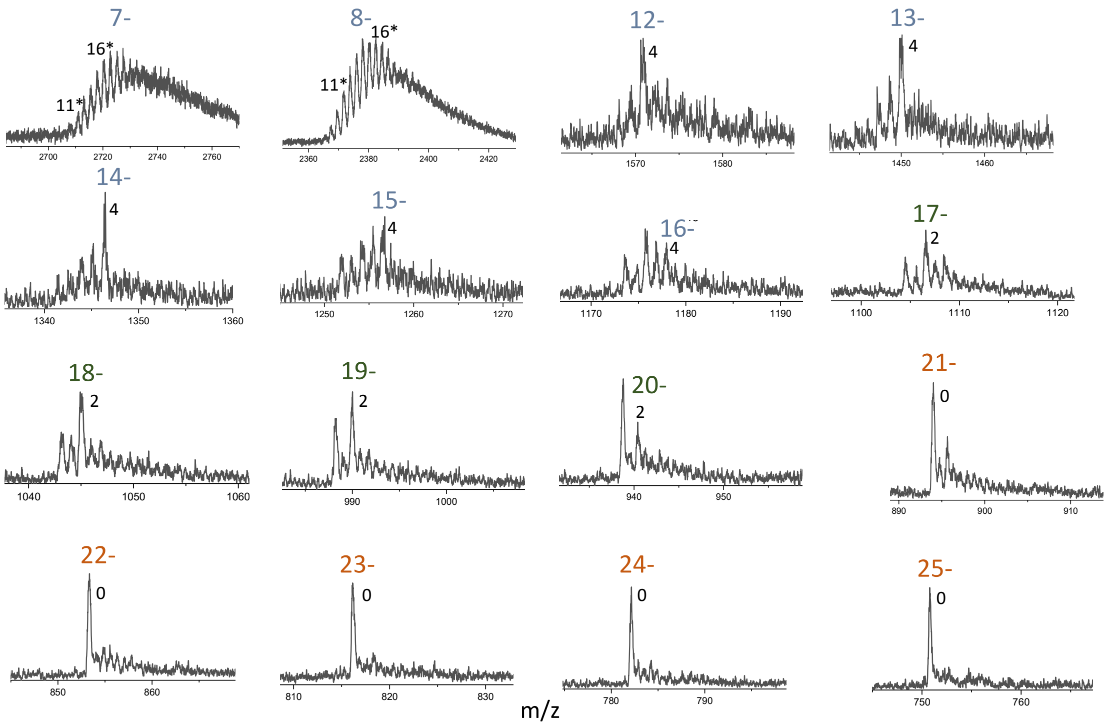


**Figure S16:** MS of G4TnG4 recorded at 320V show multiple $NH_4^+$-adducts at low charge states and very few adducts at high charge states in 50 mM $NH_4OAc$. The numbers indicate the count of non-specific $NH_4^+$-adducts (marked with an asterisk *), and specific adducts (without asterisk).

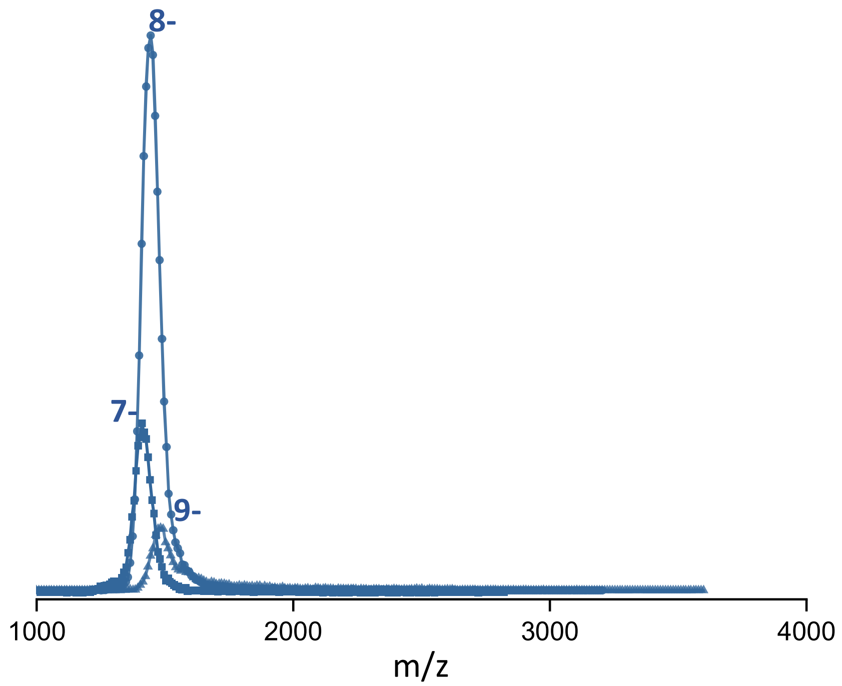


**Figure S17:** CCSD of G4TnG4 under native supercharging condition in presence of 0.4% of propylene carbonate in 150 mM aqueous $NH_4OAc$ solution showing unimodal distribution with few charge states.

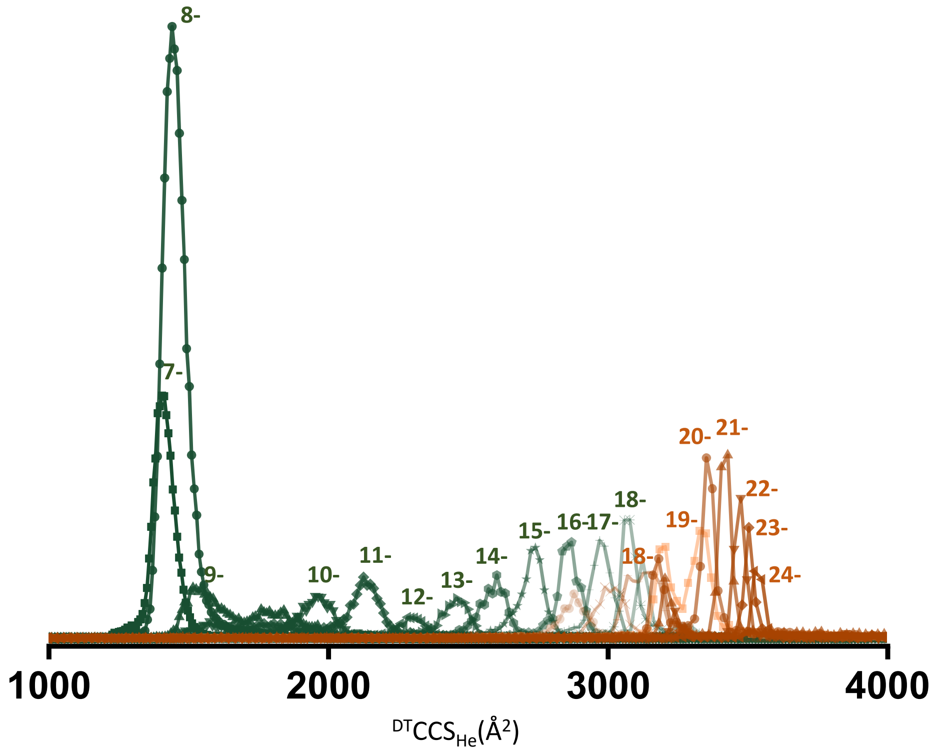


**Figure S18:** CCSD of G4Tn under native supercharging condition in presence of 0.4% of propylene carbonate in 150 mM aqueous $NH_4OAc$ solution. It exhibits a second distribution of CCS centered on 11- charge state, revealing previously hidden charge states. Under this condition, all four distributions of CCS are clearly observed.

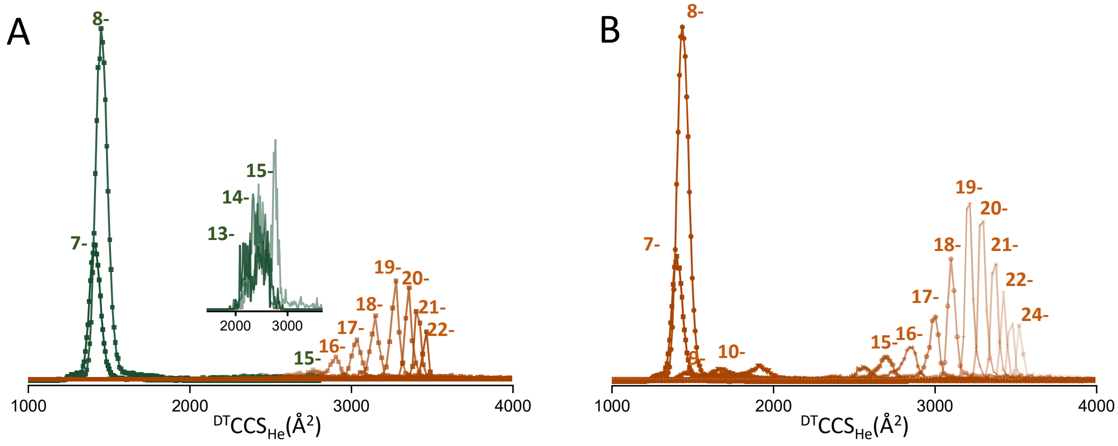


**Figure S19:** CCSD of TnG4Tn (A), NG (B), under native supercharging condition in presence of 0.4% of propylene carbonate in 150 mM aqueous $NH_4OAc$ solution. Inset in A shows the presence of 13- to 15- ions.

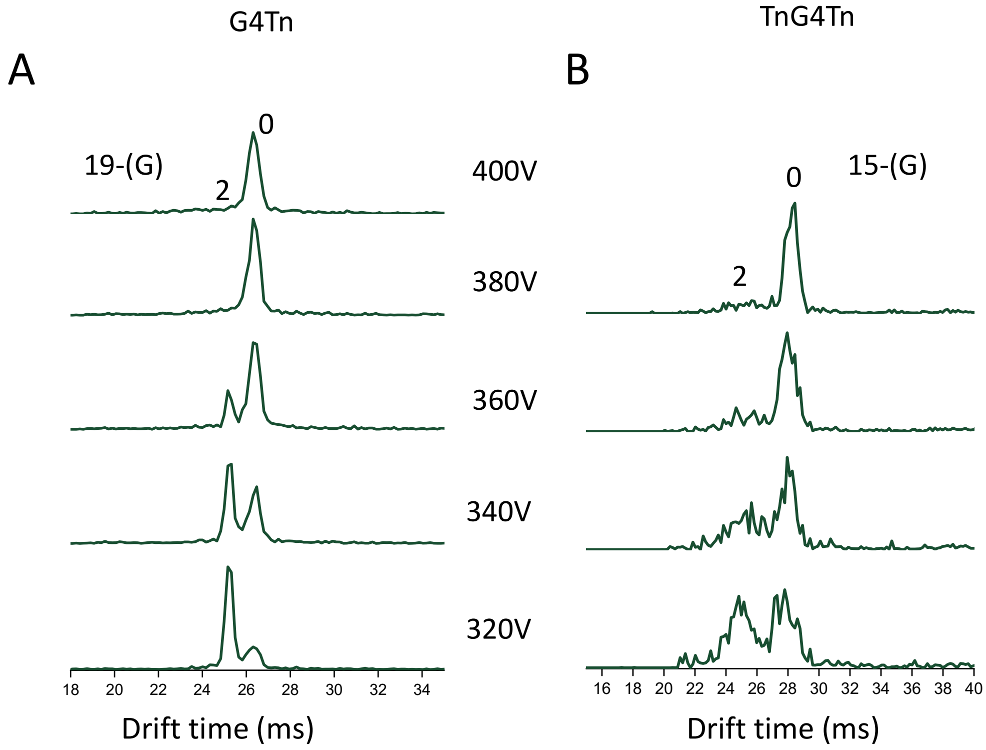


**Figure S20:** The CIU data: arrival time distributions as a function of the increasing the fragmentor voltage of G4Tn (A) and TnG4Tn (B) in 150 mM aqueous $NH_4OAc$ with 0.4% of propylene carbonate. The numbers indicate the count of specifically bound $NH_4^+$ ions. At low fragmentor voltage (320V), G4Tn predominantly adopts a folded state at 19- with two specifically bound $NH_4^+$ ions, whereas TnG4Tn displays two equally abundant distributions at 15-, one with two specifically bound $NH_4^+$ ions and another lacking $NH_4^+$ ions. The arrival time distribution plot clearly illustrates the differences in the gas-phase stability of TnG4Tn even for lower charge (15- is shown here) in comparison to G4Tn.

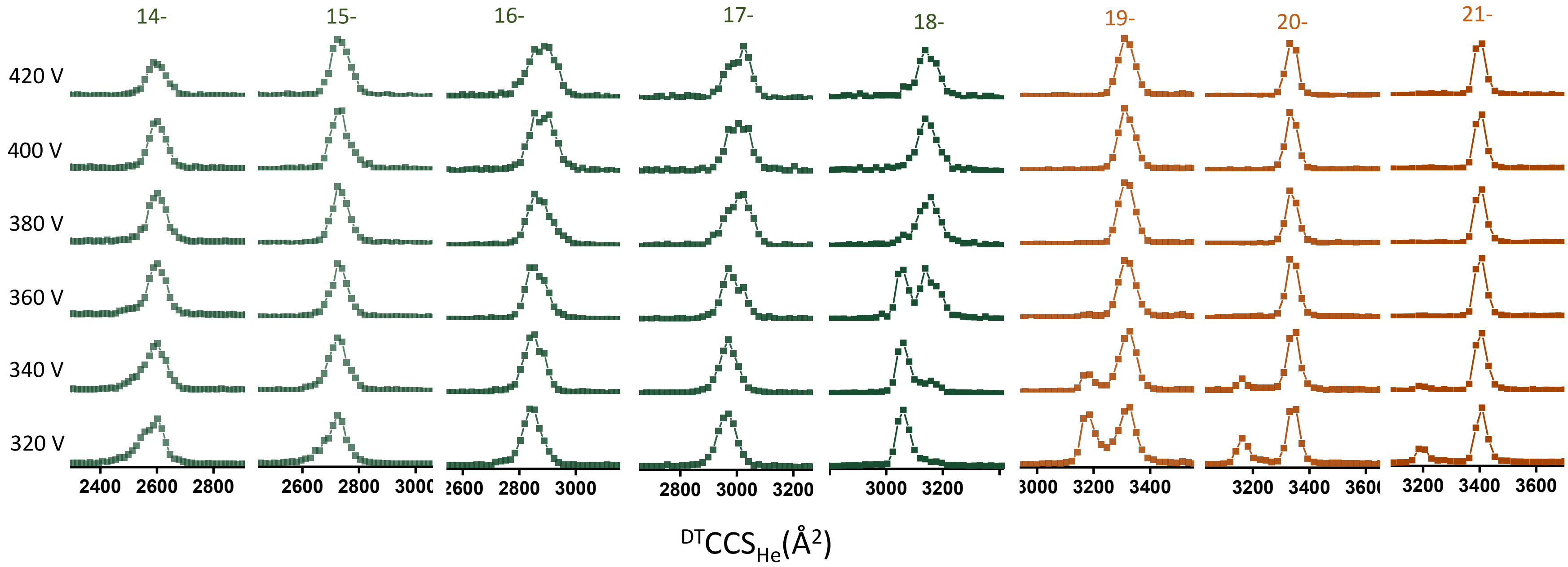


**Figure S21:** Collision-induced unfolding: evolution of the CCSDs upon increasing the fragmentor voltage (written on the left), for different charge states of G4Tn in 150 mM aqueous $NH_4OAc$ with 0.4% of propylene carbonate.

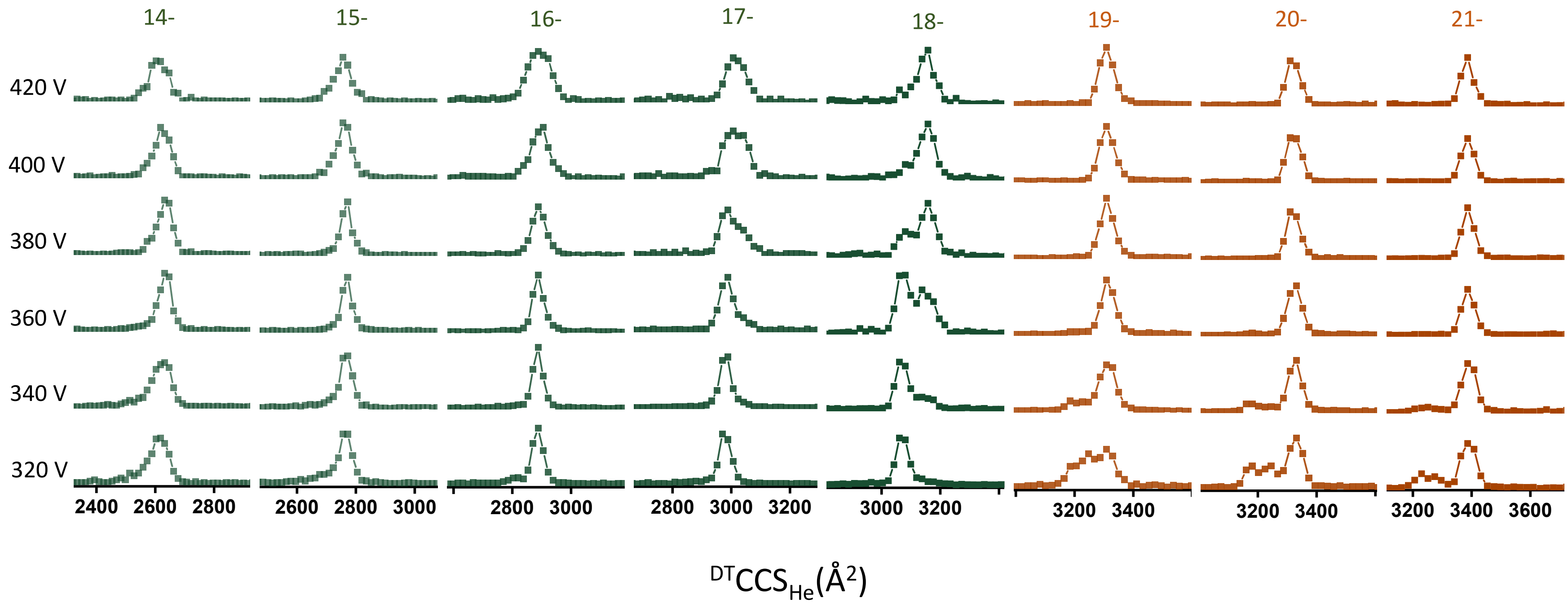


**Figure S22:** Collision-induced unfolding: evolution of the CCSDs upon increasing the fragmentor voltage (written on the left), for different charge states of TnG4 in 150 mM aqueous $NH_4OAc$ with 0.4% of propylene carbonate.

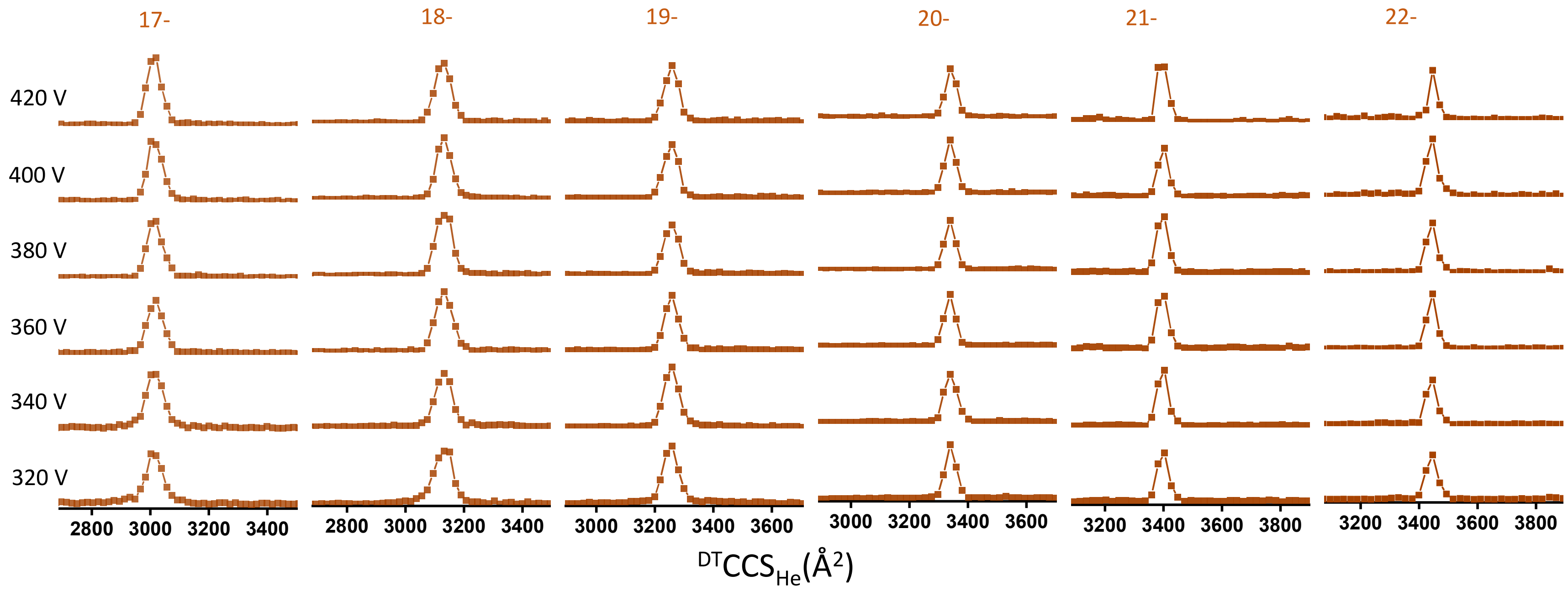


**Figure S23:** Collision-induced unfolding: evolution of the CCSDs upon increasing the fragmentor voltage (written on the left), for different charge states of TnG4Tn in 150 mM aqueous $NH_4OAc$ with 0.4% of propylene carbonate.

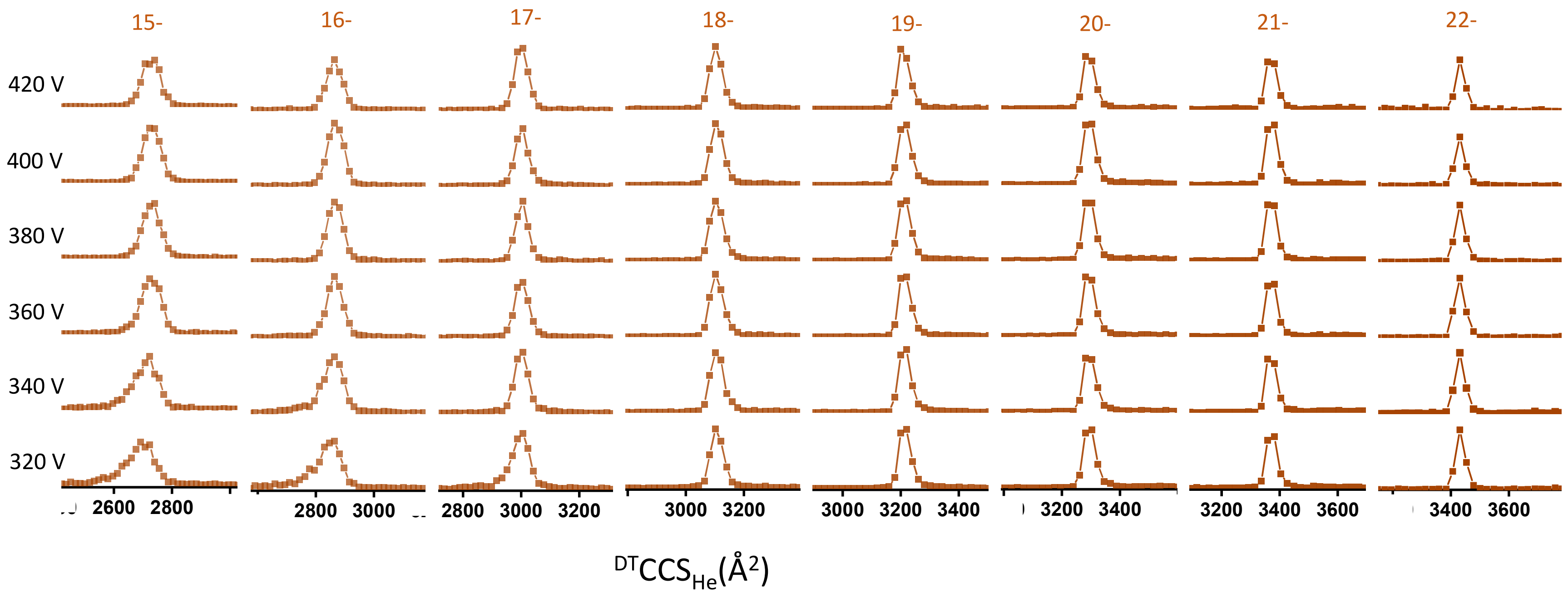


**Figure S24:** Collision-induced unfolding: evolution of the CCSDs upon increasing the fragmentor voltage (written on the left), for different charge states of NG in 150 mM aqueous $NH_4OAc$ with 0.4% of propylene carbonate.

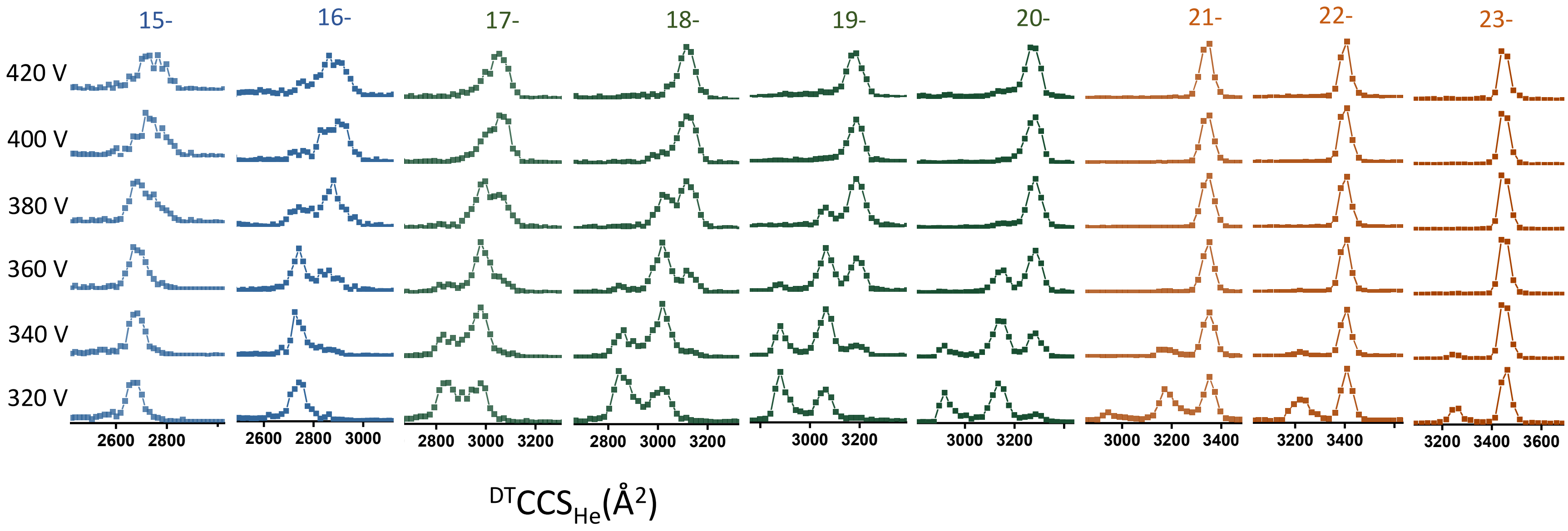


**Figure S25:** Collision-induced unfolding: evolution of the CCSDs upon increasing the fragmentor voltage (written on the left), for different charge states of G4TnG4 in 50 mM aqueous $NH_4OAc$ with 0.4% of propylene carbonate.

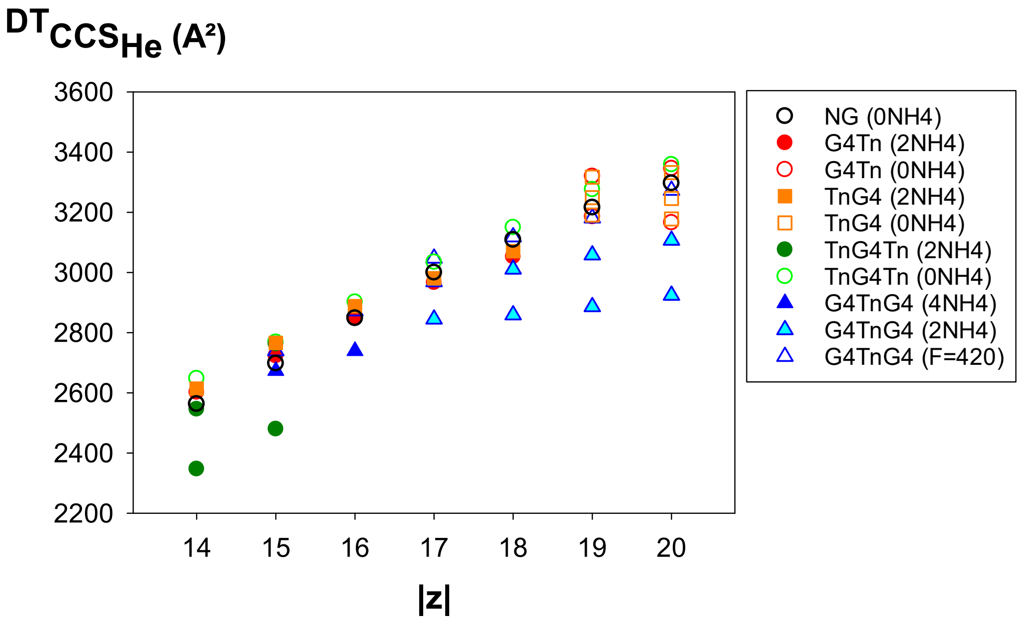


**Figure S26:** Helium collision cross sections of the 60-mer DNAs, recorded from 150 mM $NH_4OAc$ and 0.4% propylene carbonate, recorded in soft conditions (fragmentor voltage = 320 V), except for G4TnG4 which was recorded in 50 mM $NH_4OAc$ and 0.4% propylene carbonate, and for which the data of the extended forms were taken from data recorded at a fragmentor voltage of 420 V.. Filled symbols correspond to sequences with all ammonium ions and thus with preserved G-quadruplex structures.
